# Assessment of native mass spectrometry as a screening method to identify and characterize RNA-targeting small molecules

**DOI:** 10.1101/2025.02.09.637332

**Authors:** Louise M. Sternicki, Jack W. Klose, Sally-Ann Poulsen

## Abstract

The intentional targeting of RNA with small molecules is recognized as a viable pathway to new therapeutics with potential to vastly expand the proportion of the human genome considered druggable. The practical considerations to deliberately target RNA are largely under development, including optimal screening methods to identify small molecules for binding to RNA. Native mass spectrometry (nMS) is established as a valuable biophysical method for identifying small molecule hits against diverse biomolecular targets, most frequently proteins and protein-protein interactions. Herein we assess nMS for studying the binding of small molecules with RNA aptamers as a model for nMS screening of RNA as a drug target. We first develop workflows for characterizing the binding of cognate RNA aptamer ligands and then establish a nMS method to screen a small molecule library against the RNA aptamers. nMS analysis permitted identification of binders, quantitation of binding strength and generation of structure-activity relationships with some dependence on the aptamer class. This work demonstrates the utility of nMS as a complementary and efficient target-based biophysical screening method that can characterize RNA-small molecule interactions and the potential of nMS becoming a powerful enabling tool in RNA-targeting drug discovery.

## INTRODUCTION

The mechanism of most current small molecule drugs is mediated through direct targeting of proteins, with approved drugs available for just under ∼700 proteins, representing ∼0.05% of the human genome.^1^ Furthermore, only ∼1.5% of the genome encodes proteins and of this a small portion of ∼0.2% is thought to be disease related. In comparison, >50% of the human genome is transcribed into non-coding RNA which is increasingly shown to modulate cellular processes associated with disease.^1, 2^ This role has led to the supposition that intentional targeting of RNA with small molecules could expand the druggable genome by more than an order of magnitude, and in particular provide an avenue to develop therapeutics for diseases where there is currently no treatment.^2^ Furthermore, it has been suggested that RNA may be a critical target for the future of drug discovery.^3^ Proof-of-principle targeting RNA with small molecules is exemplified by the discovery and development of risdiplam, an FDA approved small molecule drug (Evrysdi, approved in August 2020) for the treatment of spinal muscular atrophy (SMA), where the mode of action is, in part, through binding to RNA.^4^ Risdiplam was however discovered through phenotypic screening and given RNA’s potential to modulate disease, there is a need to develop robust target oriented workflows for intentional and rational targeting of specific RNAs to identify small molecule binders as hit starting points for therapeutic development.^5, 6^ As RNA comprises more limited primary and secondary structural features compared to proteins, the identification of specific drug-like small molecules for RNA is regarded as more challenging than for classical protein drug targets.^3, 7^ Disney and colleagues have reviewed methods so far explored for screening small molecules against RNA targets.^3, 8^ These methods encompass affinity mass spectrometry,^9, 10^ fluorescence-based assays,^11, 12^ microarray-based screening^13^ and DNA-encoded compound library (DEL) technology,^14^ concluding that new methods are needed to support the field. Furthermore, computational approaches have also been utilized together with biophysical methods to probe RNA interactions with small molecules.^15^ The primary structure of unmodified RNA comprises four natural RNA bases, adenine (A), guanine (G), cytosine (C) and uracil (U), while RNA folds to generate three dimensional structural elements that provide potential pockets for molecular recognition and supporting specific binding of small-molecules. These elements include helical regions, hairpin loops, bulges, multibranched loops, internal loops, as well as longer-range interactions (e.g. pseudoknots), Figure 1.^3^ Longer length RNA may have opportunity to provide for more structural complexity than shorter RNAs.

**Figure 1.**
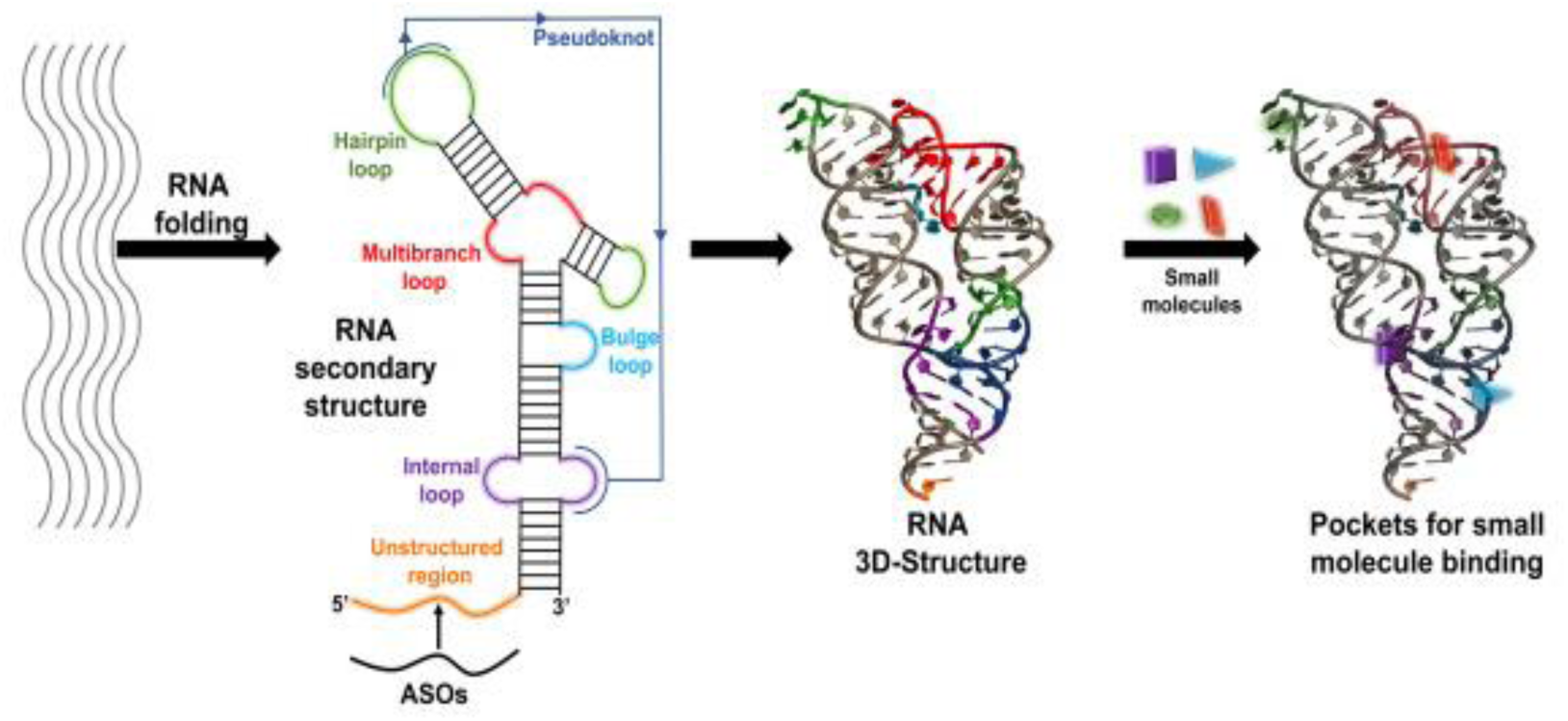
Coding or noncoding RNAs can adopt a three-dimensional structure that includes various structured elements where small molecules could specifically bind. Reprinted from SLAS Discovery. Volume 25/edition 8, Haniff HS, Knerr L, Chen JL, Disney MD, Lightfoot HL., Target-Directed Approaches for Screening Small Molecules against RNA Targets, p869-894. Copyright (2020), with permission from Elsevier.^3^

Native mass spectrometry (nMS) is established as a biophysical technique to characterize noncovalent interactions between intact, natively folded proteins and their interacting partners including small molecule ligands.^16–18^ Although nMS has been used to directly observe RNA-small molecule binding,^19–21^ the method is not routinely performed for the analysis of noncovalent interactions of RNA, especially not in the scale or context of screening compound libraries to identify novel small molecules that target RNA.^19^ Furthermore, due to the negatively charged phosphate backbone of RNA, the nature and concentration of cations may influence the structure and stability, such that when characterizing RNA using nMS the sample components and experimental instrument parameters need to be tuned to preserve signal intensity and resolution in addition to retaining a meaningful RNA structure and specific noncovalent interactions. The fundamentals of mass spectrometry for studying nucleic acid structures have been reviewed recently, concluding that nMS should become an indispensable method for advancing RNA as a druggable target.^19^

RNA aptamers are a subset of RNA types comprising single-stranded sequences of 20-40 oligonucleotides that fold into a well-defined and stable three-dimensional structure that is optimized to bind to a specific small molecule ligand (their cognate ligand) with high affinity and specificity via noncovalent interactions.^5, 6, 22^ RNA aptamers are most commonly generated using <u>S</u>ystematic <u>E</u>volution of <u>L</u>igands by <u>EX</u>ponential enrichment (SELEX), with RNA aptamers developed for diverse target classes including metal ions, small molecules and proteins as well as intact prokaryotic and eukaryotic cells.^5, 6^ Herein we investigate the scope of nMS as a biophysical method for characterizing ligand binding to four RNA aptamers as model systems for structured RNA more generally. We then employ nMS as a screening platform against RNA with a curated small molecule compound library to evaluate binding, specificity, selectivity and diversity of RNA-ligand interactions with the goal to provide a robust workflow that may be applied as a routine screening method in both industry and academic settings to identify drug-like novel RNA binding ligands.

## RESULTS AND DISCUSSION

Four RNA aptamers and their corresponding cognate ligands are employed in this study, Table 1 and Figure 2. This includes two aminoglycoside RNA aptamers (tobramycin and kanamycin B aptamers) selected as their corresponding ligands, aminoglycoside antibiotics, are a compound class with a therapeutic pedigree that is inclusive of binding to ribosomal RNA to inhibit protein synthesis.^23, 24^ The core structure of aminoglycosides consists of a 2-deoxystreptamine or streptamine moiety with amino sugar substituents at different positions.^24^ The mechanism of action is driven by the presentation of basic amino groups and hydrogen bond donor moieties that participate to provide a scaffold for a network of interactions with the ribosomal RNA targets,^23^ while it has been demonstrated that both electrostatics and conformational flexibility play a significant role in the ability of aminoglycosides to bind to RNAs more generally.^4, 25^ Of note is that the tobramycin RNA aptamer has been characterized by nMS previously for binding to several aminoglycosides^26^ and thus it provides a model RNA aptamer to benchmark against for the more extensive small molecule binding studies explored using nMS herein. As the ligand properties of aminoglycosides sit well outside of the usual rule-of-five parameters for orally bioavailable small molecule drugs (e.g. tobramycin has 14 hydrogen bond acceptors, 10 hydrogen bond donors and log P -2.8),^27^ we also selected two purine RNA aptamers (xanthine and theophylline aptamers) as we were interested in RNA that bound to compounds that were more representative of screening hits in drug discovery campaigns. The structures of purines have contrasting properties to aminoglycosides, they comprise a flat heteroaromatic core commonly with simple substituents and no or limited stereocentres, less hydrogen bond donors and acceptors, and overall lower complexity structures. Although xanthine and theophylline are not therapeutics, they do exhibit bioactivity as important signalling molecules, and they are also considered as fragments (MW <300 Da) in the context of fragment-based drug discovery. This renders them of particular interest because fragments are proven optimal starting points for successful drug discovery outcomes with so far only a few fragment studies with RNA.^28–30^

**Table 1.** Selected properties of RNA aptamers used in this study.

| RNA Aptamer | Sequence | Length | Molecular weight (Da) |
| --- | --- | --- | --- |
| Tobramycin | 5'-G<sub>3</sub>ACU<sub>2</sub>G<sub>2</sub>U<sub>3</sub>AG<sub>2</sub>UA<sub>2</sub>UGAGUC<sub>3</sub>U-3' | 27-mer | 8,670 |
| Kanamycin B | 5'-G<sub>3</sub>AGCUCG<sub>2</sub>UAC<sub>2</sub>GA<sub>2</sub>U<sub>2</sub>CUC-3' | 22-mer | 7,030 |
| Xanthine | 5'-G<sub>2</sub>CACGUGUAU<sub>2</sub>ACCCUAGUG<sub>2</sub>UCGACGUGC<sub>2</sub>-3' | 32-mer | 10,230 |
| Theophylline | 5'-A<sub>2</sub>GUGAUAC<sub>2</sub>AGCAUCGUCU<sub>2</sub>GAUGC<sub>3</sub>U<sub>2</sub>G<sub>2</sub>CAGCACU<sub>2</sub>CA-3' | 42-mer | 13,370 |

**Figure 2.**
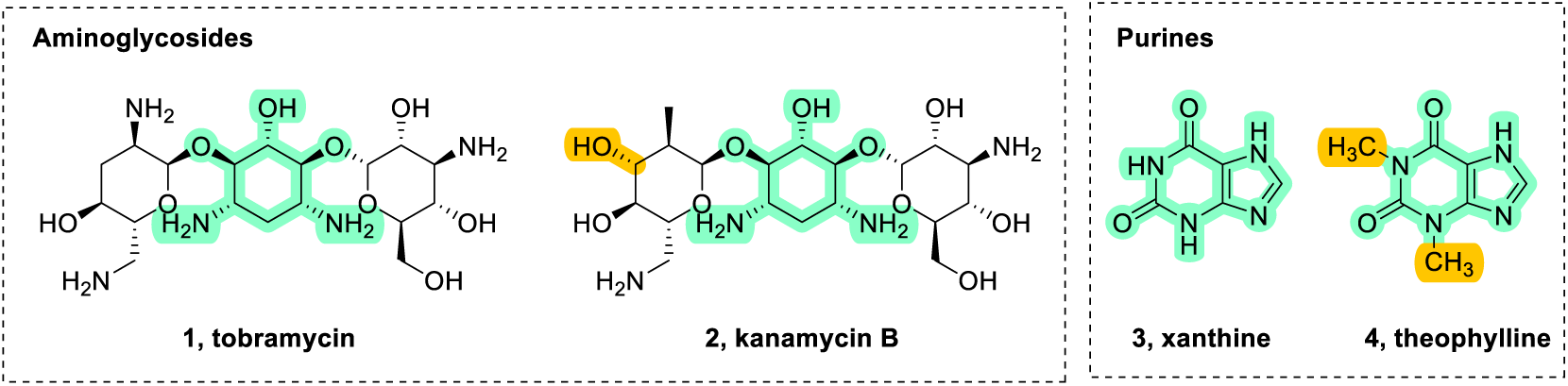
Structure of the cognate ligands used to generate the corresponding RNA aptamers of Table 1. Aminoglycoside ligands tobramycin **1** and kanamycin B **2**; purine ligands xanthine **3** and theophylline **4**. The core structures are highlighted in green and key difference in structures is highlighted in orange. A panel of additional aminoglycosides (**5-8**) were included in our study as some RNA aptamer binding information is available for these ligands and, therefore, they could enable a more comprehensive assessment of the utility of nMS for characterizing RNA ligand binding and selectivity for comparison with the application of nMS with a larger screening campaign, Figure 3.

**Figure 3.**
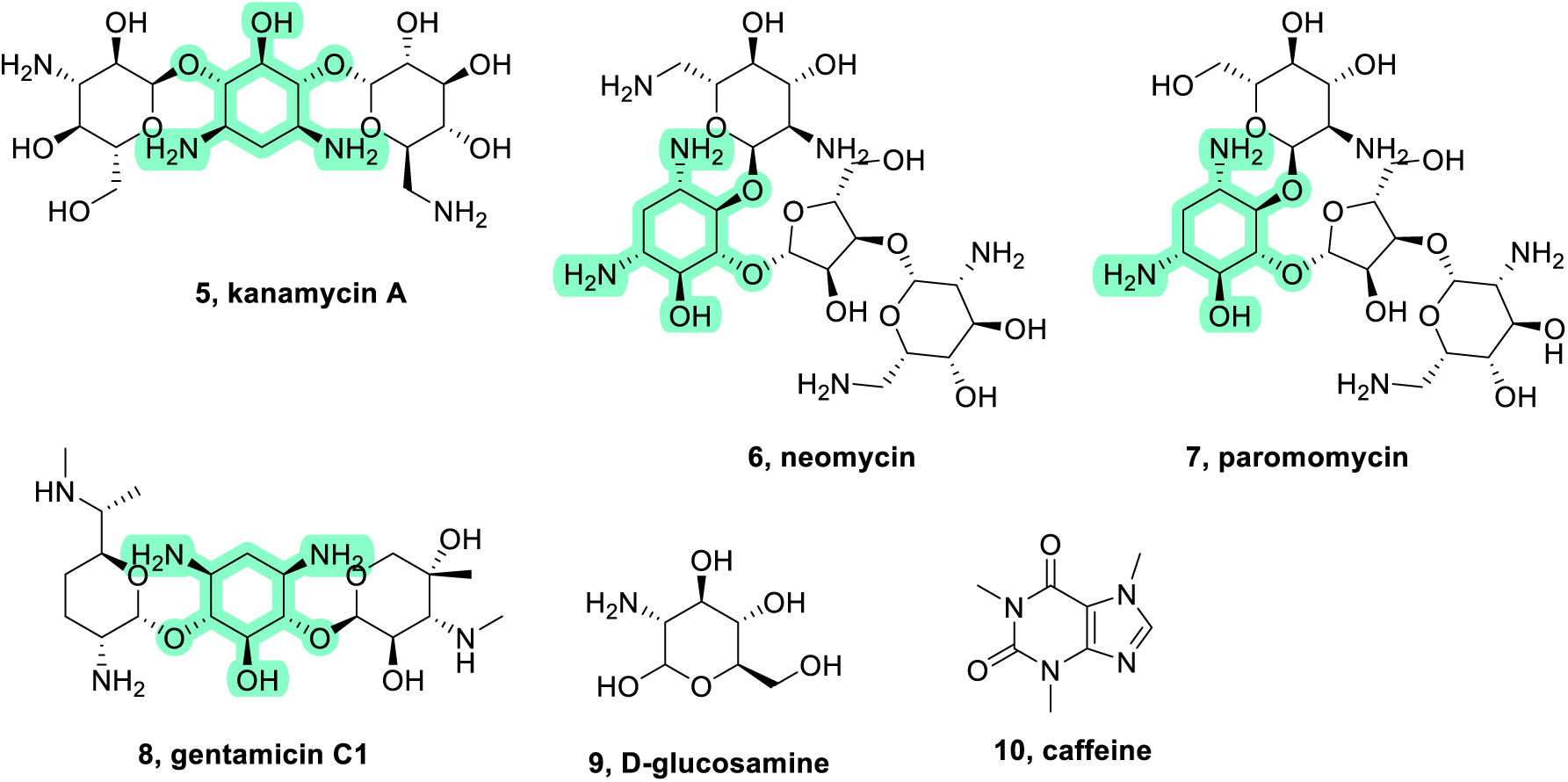
Structures of control ligands comprising additional aminoglycosides (**5-8**) and control compounds D-glucosamine (**9**) (negative control for aminoglycoside RNA aptamers) and caffeine (**10**) (negative control for purine RNA aptamers). The core structure of the aminoglycoside ligands is highlighted in green.

The tobramycin RNA aptamer sequence used in our study is the minimal X1sl aptamer, Table 1. It has two reported binding sites for the tobramycin ligand **1**, the first is high affinity with a *K*_D_ of 12 ± 5 nM, while the second is lower affinity with a *K*_D_ of 13,200 ± 500 nM, as measured using a steady-state fluorescence assay with a fluorescent tobramycin derivative, Table 2.^31^ Also reported is qualitative aptamer binding to aminoglycosides that differ in structure to tobramycin by varied hydrogen, hydroxyl and amino group substitutions on the sugar moieties, with binding of kanamycin B **2** similar to that of tobramycin and binding of kanamycin A **5** and gentamicin **8** (the fewest number of primary amino groups) weak, Table 2.^32^ There is a NMR solution structure of the aptamer bound to tobramycin **1** (PDB:2TOB) showing the aptamer with a distorted hairpin structure (six base-pair stem and a 14-residue loop region). The structure provides a binding pocket and potential network of intra/intermolecular hydrogen bonds for high affinity ligand binding to tobramycin **1** with features discriminating binding for other aminoglycosides.^32^ A full length tobramycin aptamer (W13 aptamer, 109-mer) is reported to have lower binding affinity for tobramycin **1** (*K*_D_ = 4,300 ± 700 nM), similar affinity for neomycin **6** (*K*_D_ = 4,700 ± 900 nM) and 10-fold weaker affinity for gentamicin **8** (K_D_ = 47,000 ± 11,000 nM), Table 2.^31^ D-glucosamine **9**, a simple amino sugar, was used for counterselection in the development of the tobramycin RNA aptamer and is reported to not bind to this full length aptamer.^31^ Binding of a full-length kanamycin B RNA aptamer (110-mer, K8) with kanamycin B **2** is reported with a *K*_D_ of 181 nM, and, interestingly, even though selected against kanamycin B **2** the reported binding affinity with tobramycin **1** is 10-fold higher (*K*_D_ of 11.6 nM) as measured using fluorescence anisotropy, Table 2.^33^ Binding to other aminoglycosides was weaker (kanamycin A **5**, *K*_D_ of 4.4 µM; neomycin **6**, *K*_D_ of 1.1 µM; paromomycin **7**, *K*_D_ of 1.5 µM), Table 2.^33^ There is no binding data or high resolution structure available for the minimal kanamycin B RNA aptamer (22-mer, K8-1-1) used in our study, however, computational and aptamer truncation and footprinting experiments predict that the aptamer forms a stem-loop structure and pockets for potential binding of ligands.^33^ In summary, the reported specificity of binding to structurally related ligands reported for the aminoglycoside aptamers ranges from strong to very weak binding, with ligand structure-activity relationships likely correlating with both the potential for electrostatic interactions including hydrogen bonding or steric interactions related to the ligand and/or aptamer structure.

**Table 2.** Reported aminoglycoside binding (qualitative) or binding affinity (*K*_D_ in nM) to the minimal tobramycin RNA aptamer (X1sl) used in this study, and comparison binding with a full-length tobramycin RNA aptamer (W13) and full-length kanamycin B RNA aptamer (K8).^26, 31–33^

| Compound | Ligand | MW<br>(Da) | Tobramycin<br>aptamer X1sl | Tobramycin<br>aptamer W13 | Kanamycin B<br>aptamer K8 |
| --- | --- | --- | --- | --- | --- |
| **1** | tobramycin | 467.52 | 12 ± 5 (site 1)<br>13200 ± 500 (site 2) | 4300 ± 700 | 11.6 |
| **2** | kanamycin B | 483.51 | similar to tobramycin | <sup>b</sup> | 181 |
| **5** | kanamycin A | 484.50 | weak binding | <sup>b</sup> | 4,400 |
| **6** | neomycin | 614.65 | <sup>b</sup> | 4700 ± 900 | 1,100 |
| **7** | paromomycin | 615.63 | <sup>b</sup> | <sup>b</sup> | 1,500 |
| **8** | gentamicin | 477.60 | weak binding | 47000 ± 11000 | <sup>b</sup> |
| **9** | D-glucosamine<sup>a</sup> | 179.17 | <sup>b</sup> | no binding | <sup>b</sup> |

<sup>a</sup>Counter selection ligand (simple amino sugar) for tobramycin RNA aptamers.<sup>31</sup> <sup>b</sup>No reported binding data.

The xanthine RNA aptamer (32-mer) used in our study was developed against xanthine **3** (MW 152.11 Da) covalently linked to an agarose column at the C8 position of **3** and is suggested to form a stem-loop structure with an internal loop, similarly to the kanamycin B aptamer. The dissociation constant for xanthine **3** was determined using an equilibrium filtration procedure with tritium-labelled ligands and is reported with a *K*_D_ of 3.3 ± 0.2 µM.^34^ The *K*_D_ values for various xanthine analogues were also determined, revealing that the N1H, N7 and O6 of **3** are likely involved in the binding with the RNA aptamer and that guanine (with an amino group in place of an exocyclic carbonyl) has an affinity higher than xanthine (*K*_D_ of 1.3 ± 0.1 µM).^34^ Lastly, the theophylline RNA aptamer used in our study is reported with a binding affinity *K*_D_ of 0.32 ± 0.13 µM for theophylline **4** (MW 180.16 Da) with an absolute requirement for at least 5 mM Mg^2+^, Co^2+^ or Mn^2+^ cations (in the absence of these cations theophylline **4** binding has a *K*_D_ of ∼ 4 mM).^35^ No binding with the aptamer is observed for **4** with other divalent or lanthanide cations indicating that Mg/Co/Mn may be structural cations in the core of the theophylline RNA aptamer while the diffuse cation requirement can be met by various different cations.^35, 36^ A pH above 6.5 was required to facilitate correct base pairing of the aptamer backbone (prevent C27-A7H+ base pairing) and to structure the aptamer for theophylline binding, with the aptamer forming a stem-loop structure with an internal loop and mismatched bulge.^35^ A team at Novartis Institute for BioMedical Research reported a proof-of-concept high throughput screen of ∼1 million compounds (MW <600 Da, 50 µM) against fluorescently labelled theophylline aptamer using a strand invasion assay.^7^ A 0.014% hit rate was reported, with 46 hits identified with affinity greater than theophylline **4**. The assay buffer contained 150 mM NaCl, 5 mM MgCl_2_, 10 mM HEPES (pH 7.4), and 0.01–0.1% DMSO. Atomic-resolution X-ray crystal structures were obtained for the aptamer with theophylline **4** and four of the best screening hits. The X-ray crystallography structure of the theophylline **4** bound aptamer revealed a structural role for three bound cations, although they did not coordinate to **4** directly.^7^ It has been shown that RNA crystallisation conditions can affect the position and occupancy of metal ions even for the same RNA molecule and also that ion occupancy in solution can differ from that in the crystal structure.^37^

The SELEX selection conditions for development of RNA aptamers use high salt concentrations (140-500 mM NaCl, 1-5 mM MgCl_2_ plus other salt additives for the aptamers in this study).^31, 33, 34, 36^ Of relevance to nMS is that nonspecific mono- and divalent cations may be essential to support RNA folding (e.g. demonstrated as critical for the theophylline aptamer ^35^) influencing RNA structure, stability and noncovalent interactions through charge screening of the negatively charged RNA phosphate groups.^19, 37^ These salts are, however, nonvolatile and thus generally unsuitable for nMS samples as they would lead to extensive salt adducts during the electrospray process and ion suppression, significantly degrading native mass spectra quality.^19, 38^ For nMS analysis, ammonium acetate (NH_4_OAc) solutions are commonly used in nMS samples of biomolecules to provide the needed ionic strength (e.g., 150 mM NH_4_OAc mimics the physiological ionic strength) and pH to maintain a native structure and specific noncovalent interactions but with good mass spectra quality, for strongly charged RNA this is a particularly important consideration.^19, 37–39^ It has been reported that ammonium ions (NH_4+_) can stabilize RNA to an extent similar to sodium ions (Na^+^),^19^ and that the relevant (qualitative) charge screening of divalent cations is 10-fold greater than monovalent cations.^37^ In mammalian cells, where eventual therapeutics that target RNA would have their effect, typical concentrations of free Na^+^ is 10 mM while free Mg^2+^ is 1 mM.^37^ NH_4+_ ions, which ultimately generate protons in the electrospray process,^39^ however may not solely be able to replace or account for specific structural cations if essential for RNA folding or binding. Optimising ion transfer in nMS thus requires instrument parameters to remove extensive non-structural salt adducts while maintaining native RNA structure and any specific noncovalent interactions whilst mitigating the potential for nonspecific interactions.

The first reported nMS study of RNA aptamers binding to their small molecule ligand partners described two RNA aptamers, the tobramycin (same as our study, 22-mer minimal X1sl aptamer, Table 2) and flavin mononucleotide (FMN, a 35-mer) RNA aptamers, with the nMS test samples comprising 2 µM aptamer with 1 equiv ligand in 50 mM NH_4_OAc in 30% isopropanol/water, 2-5 equiv divalent metal ions, pH 6.5, measured in the negative ion mode.^26^ The reported solution binding data for the tobramycin aptamer correlated well with the nMS data, with binding of tobramycin **1** and two related aminoglycoside compounds (kanamycin B **2** and gentamicin **8**) assessed.^26^ Both tobramycin **1** and kanamycin B **2** bound similarly, almost saturating a first high affinity binding site and then occupying a small amount of a second weaker affinity binding site. A titration of tobramycin **1** (1-5 equiv) displayed increasing binding to and then saturation of the first binding site and an increase in binding to the second site, however the second binding site did not saturate.^26^ In comparison, gentamicin **8** weakly bound the tobramycin RNA aptamer only partially occupying the first binding site. For the FMN aptamer, the observed nMS binding was weaker than solution phase measurements predicted.^26^ It was proposed that a low stability gas phase aptamer-ligand complex may be responsible, associated with coulombic repulsion effects between the aptamer and the FMN ligand as both aptamer and ligand are anionic.^26^ A more recent study using nMS investigating an alternate tobramycin RNA aptamer (minimal J6sl aptamer, a 27-mer) (0.5 µM, 100 mM NH_4_OAc) with tobramycin **1** (0.5-2 equiv), also in the negative ion mode, provided similar findings, with the first binding site almost saturated and weak binding to a second binding site observed.^40^ The nMS measured *K*_D_ for the first binding site (7.8 ± 1.8 nM and 11.1 ± 3.4 nM for the 6- and 5-charge states respectively) agreed with previous literature (9 nM), however, the nMS measured *K*_D_ for the second binding site (0.25 ± 0.09 µM and 0.18 ± 0.07 µM for the 6- and 5-charge states respectively) was 10-fold stronger than previous literature (2.7 µM).^40, 41^

### nMS analysis of RNA aptamers

A mass spectrum of each aptamer (2 µM, 250 mM NH_4_OAc, ±20 µM Mg(OAc)_2_, ±1% (v/v) DMSO, pH 6.5) was acquired using nMS in both the positive ion and negative ion mode. The concentration of NH_4_OAc was selected to provide sufficient ionic strength and charge screening for RNA, while ±Mg(OAc)_2_ (10 equiv) was assessed as the Mg^2+^ divalent cation had been used with the SELEX selection conditions for the aptamers, and although not observed in reported structural studies, the impact of this cation on folding and/or structure of the aptamer and for specific noncovalent interactions mediated by Mg^2+^ with their cognate ligands was of interest. Additionally, 1% DMSO (v/v) was added to each aptamer as DMSO is the solvent most used for compound library storage and thus is a control for eventual compound screening using nMS. All aptamers were visible with nMS with a narrow charge state distribution of predominantly one or two charge states. The nMS measured MW for the tobramycin, xanthine, and theophylline RNA aptamers (∼8,666-8,667 Da, ∼10,231.5 Da and 13,366-13,368 Da, respectively) corresponded with the MW expected based on the aptamer sequence, while the measured MW for the kanamycin B RNA aptamer (14,066 Da) was twice that expected based on the aptamer sequence, indicating it exists as a dimeric complex that is retained in the gas phase, Table 3. There is no reported literature with the molecular structure of this aptamer characterized and our attempts to disrupt this dimer are described later. The presence of Mg^2+^ or DMSO in the samples had minimal impact on the spectra acquired (discussed for each aptamer next). In all acquired native mass spectra the charge states observed were heavily adducted (more so with the addition of Mg^2+^ to the samples), and commonly observed with adduction of 1 to 8 Na^+^ or Mg^2+^ ions, consistent with cation-phosphate interactions that may form during desolvation.^42^ There were no significant differences in sensitivity of cognate ligand binding observed between positive and negative ion mode. A summary of observed charge states and measured MWs of the RNA aptamers are presented in Table 3.

**Table 3.** Observed charge states and measured MWs for the four RNA aptamers studied under different sample conditions (± Mg^2+^ and ± DMSO).

| RNA Aptamer | Expected MW based on sequence | Positive Ion nMS Observations |  |  |  | Negative Ion nMS Observations |  |  |  |
| --- | --- | --- | --- | --- | --- | --- | --- | --- | --- |
| | | 250 mM $\text{NH}_4\text{OAc}$ | + 20 $\mu\text{M}$ $\text{Mg}^{2+}$ | + 1% DMSO | + 20 $\mu\text{M}$ $\text{Mg}^{2+}$ + 1 % DMSO | 250 mM $\text{NH}_4\text{OAc}$ | + 20 $\mu\text{M}$ $\text{Mg}^{2+}$ | + 1% DMSO | + 20 $\mu\text{M}$ $\text{Mg}^{2+}$ + 1 % DMSO |
| Tobramycin | 8,670 Da | 4+ (major)<br>5+ | 4+ | 3+<br>4+ (major) | 4+ | 4-5- (major) | 5- | 4- (major)<br>(7- dimer) | 5- |
|  |  | 8,667.3 Da | 8,667.2 Da | 8,667.3 Da | 8,667.3 Da | 8,666.2 Da | 8,666.2 Da | 8,666.1 Da | 8,666.2 Da |
| Kanamycin B | 7,030 Da | 5+<br>6+<br>(equal intensities) | 5+<br>6+<br>(equal intensities) | 5+<br>14,066.1 Da | 5+<br>14,066.1 Da | 6-14,066.0 Da | 6-14,066.0 Da | 6-14,066.1 Da | 6-14,066.0 Da |
|  |  | 14,066.1 Da | 14,066.0 Da |  |  |  |  |  |  |
| Xanthine | 10,230 Da | 4+<br>10,231.5 Da | 4+<br>10,231.5 Da | 4+<br>10,231.5 Da | 4+<br>10,231.6 Da | 5-6-<br>(equal intensities) | 5-6-<br>(equal intensities) | 5-6-<br>(equal intensities) | 5-6-<br>(equal intensities) |
|  |  |  |  |  |  | 10,231.4 Da | 10,231.5 Da | 10,231.4 Da | 10,231.5 Da |
| Theophylline | 13,370 Da |  |  | 5+ (major)<br>6+<br>13,368.0 Da | 4+<br>5+ (major)<br>6+<br>13,366.1 Da |  |  | 6-13,365.9 Da | 6-13,366.9 Da |

#### Tobramycin aptamer

The tobramycin RNA aptamer was visible in the positive ion mode predominantly as the 4+ charge state, and in the negative ion mode in predominantly the 5-charge state, Figure 4. The addition of Mg^2+^ did not significantly change the observed charge states in either the positive or negative ion mode but increased adduct levels. The addition of tobramycin **1** (1 equiv) saturated the aptamer in both ion modes with and without Mg^2+^ present, suggesting Mg^2+^ is not essential for cognate ligand binding. Additionally, a small amount of binding of a second tobramycin ligand was observed, consistent with the literature where a second weaker affinity binding site of the tobramycin RNA aptamer for aminoglycosides is reported.^31^ The charge state distribution with DMSO depended on other sample components (ligand **1** and Mg^2+^), and was either unchanged or lowered, with lowering consistent with the effect of low concentrations of DMSO on nMS of proteins where the shift to lower charge states is attributed to slight compaction of the protein.^43^

**Figure 4.**
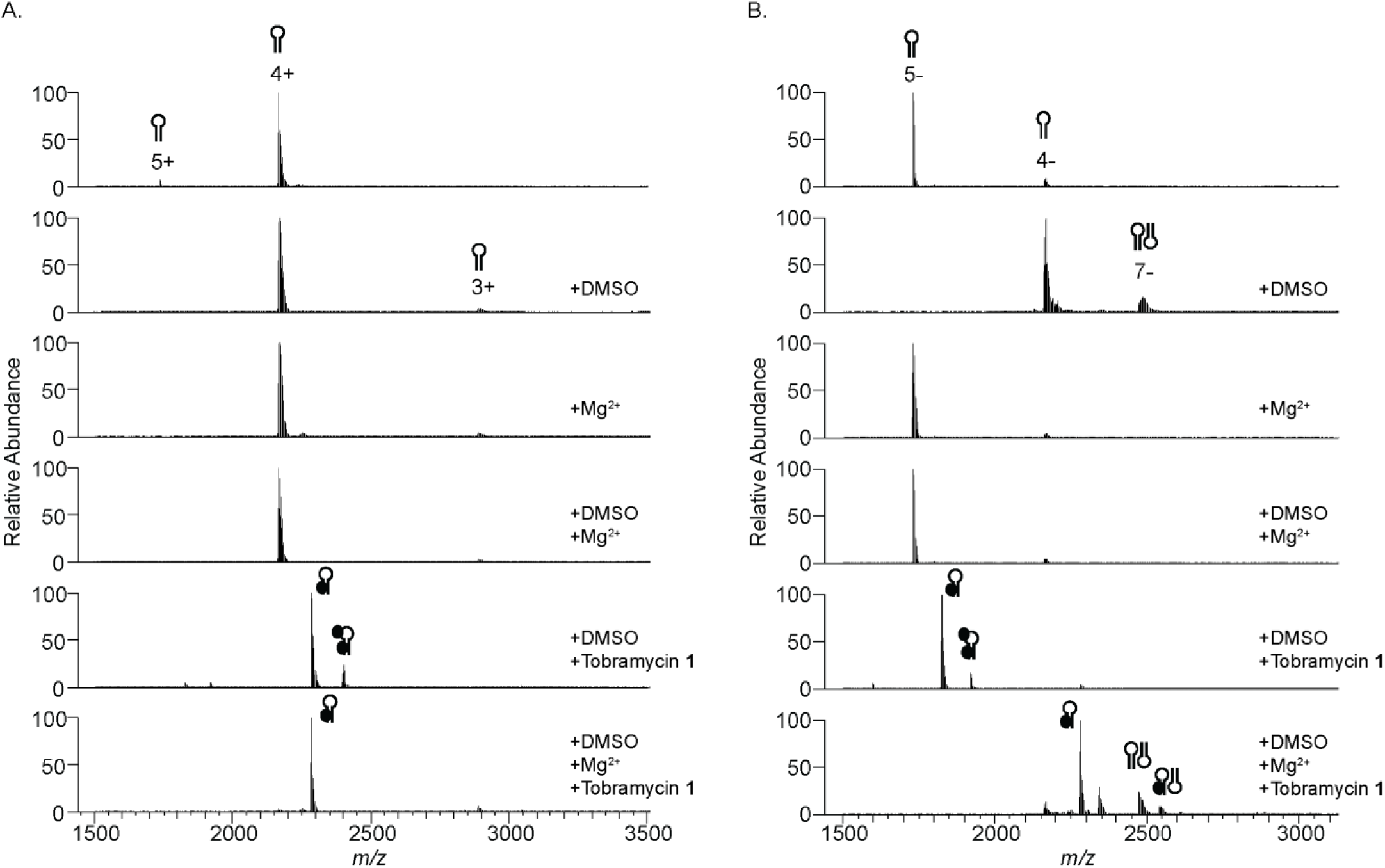
nMS analysis of tobramycin RNA aptamer (2 µM, 250 mM NH_4_OAc, pH 6.5) ± 20 µM Mg(OAc)_2_, ± 1% DMSO, ± tobramycin ligand **1** (2 µM) in (a) positive and (b) negative ion mode. Spectrum annotations show sample conditions. Hairpin represents aptamer, black circle represents bound ligand **1**.

#### Kanamycin B aptamer

The kanamycin B RNA aptamer was visible as a dimer with 5+ and 6+ charge states of equal intensity in the positive ion mode and predominantly as 6-charge state in the negative ion mode, Figure 5. The addition of Mg^2+^ did not change the observed charge states in either mode, while DMSO did not change the observed charge states in negative ion mode but in the positive ion mode the 6+ charge state was substantially diminished with only the 5+ charge state predominantly observed. Attempts to disrupt the kanamycin B dimer using solvents (denaturants, acid, base, higher amounts of DMSO) or refolding steps (both pre and post buffer exchange) to measure only the monomer were not successful (data not shown). The cognate ligand kanamycin B **2** (1 equiv) saturated the dimeric aptamer in both the positive and negative ion modes, with a small amount observed with two ligands of **2** bound. This may suggest the occupancy of one ligand per aptamer RNA strand or binding of the ligand at alternate sites such as within the dimer interface. Similarly to the tobramycin aptamer, Mg^2+^ did not alter the mass spectrum of the kanamycin B aptamer with its cognate ligand, suggesting this cation is not required for ligand binding.

**Figure 5.**
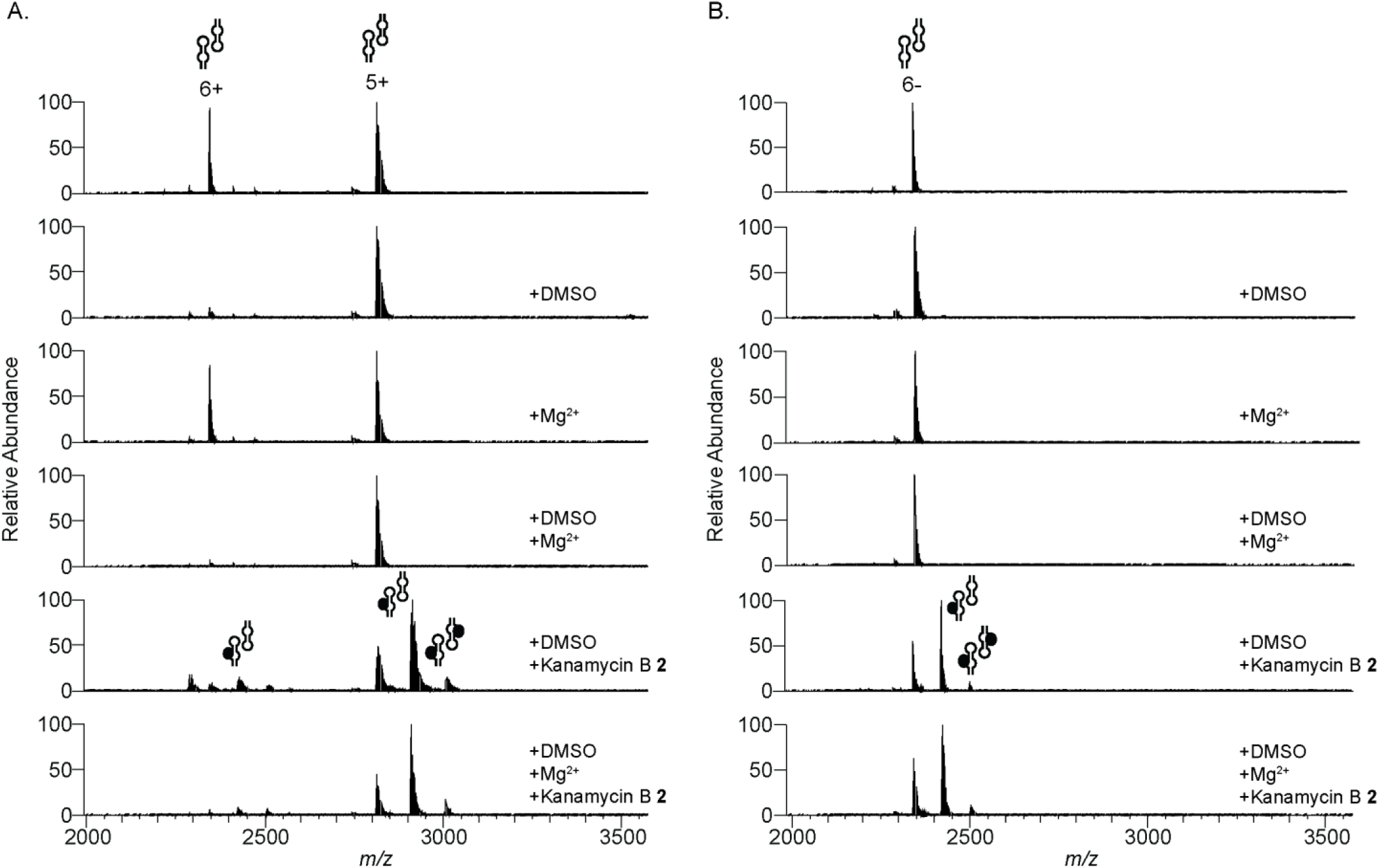
nMS analysis of kanamycin B RNA aptamer (2 µM, 250 mM NH_4_OAc, pH 6.5) ± 20 µM Mg(OAc)_2_, ± 1% DMSO, ± kanamycin B ligand **2** (2 µM) in (a) positive and (b) negative ion mode. Spectrum annotations show sample conditions. Twin hairpin represents dimeric aptamer, black circle represents bound ligand **2**.

#### Xanthine aptamer

The xanthine RNA aptamer was visible in the positive ion mode as a 4+ charge state species and in the negative ion mode as a 5- and 6-charged states species of similar intensity. The addition of DMSO or Mg^2+^ did not change the observed distribution of charge states, Figure 6. Binding of the cognate ligand xanthine **3** (1 equiv, then 10 equiv) was not detected in either the positive or negative ion modes, with or without Mg^2+^, when using the same MS instrument parameters as used with the aminoglycoside RNA aptamers above. As the reported *K*_D_ for xanthine **3** binding to its aptamer is an order of magnitude lower than the *K*_D_ of **1** and **2** with their corresponding RNA aptamers, we considered exploring gentler MS instrument parameters: source DC offset lowered (21V →10 V), capillary temperature lowered (275 °C → 100 °C), extended trapping slightly increased (1 eV → 1.2 eV), trapping gas pressure lowered (1.0 → 0.5). With these modified parameters, binding of xanthine **3** (10 equiv) to the aptamer was observed with a minor amount of a second xanthine **3** ligand bound, however, the nMS spectra had heavily increased adduct levels which lowered the overall sensitivity of the nMS analysis and lessened the utility of applying gentler instrument parameters, Figure 6(c).

**Figure 6.**
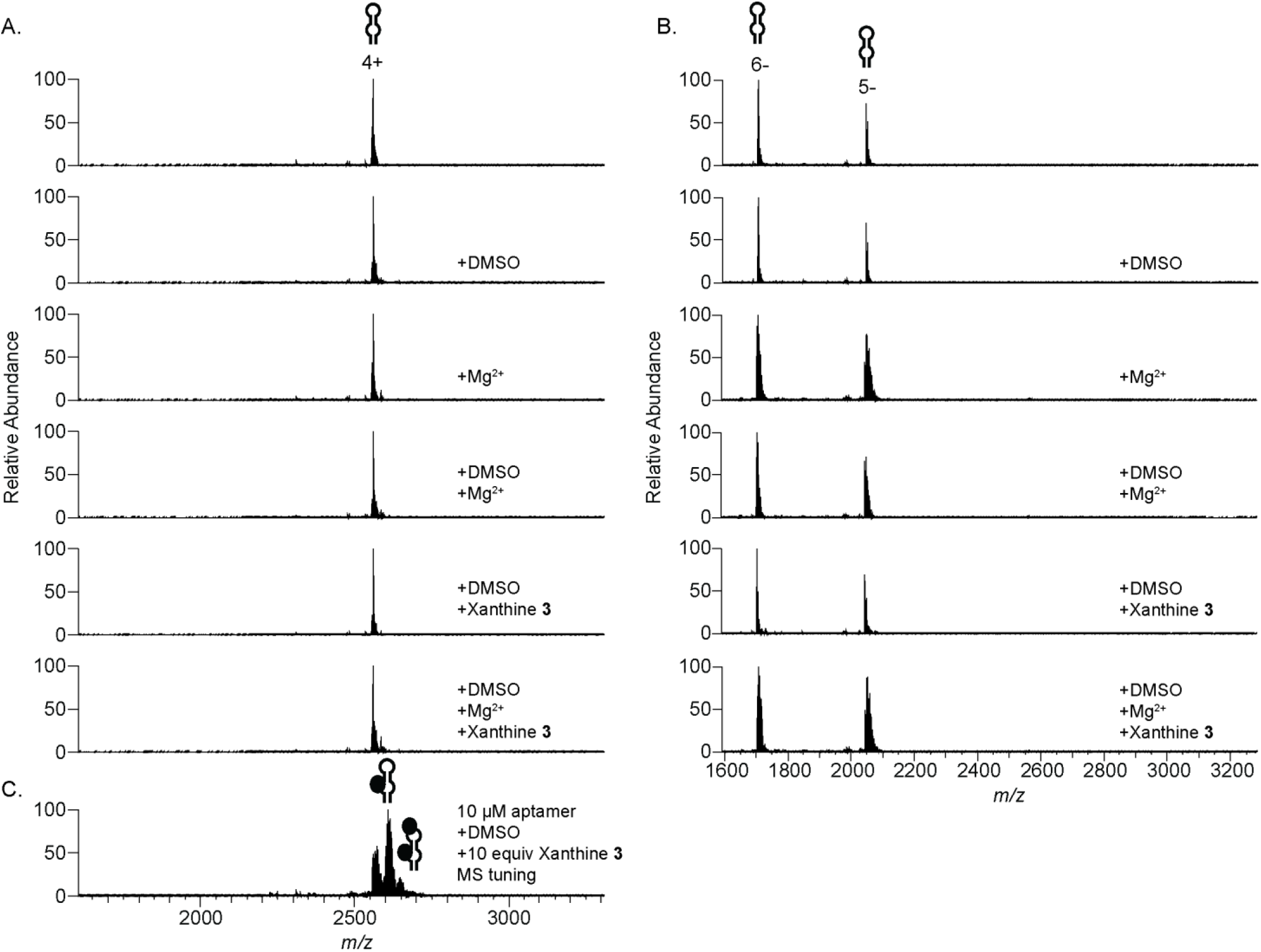
nMS analysis of xanthine RNA aptamer (2 µM, 250 mM NH_4_OAc, pH 6.5) ± 20 µM Mg(OAc)_2_, ± 1% DMSO, ± xanthine ligand **3** in (a) positive and (b) negative ion mode, and (c) with gentler MS instrument parameters required to observe **3** binding. Spectrum annotations show sample conditions. Hairpin represents aptamer, black circle represents bound ligand **3**.

#### Theophylline aptamer

The theophylline RNA aptamer (1% DMSO, with and without Mg^2+^) was visible in the positive ion mode with 5+ as the predominant charge state (4+ and 6+ also observed) and in the negative ion mode with 6-the sole charge state, Figure 7. With addition of theophylline **4** (up to 10 equiv), no ligand binding was observed in either ion mode. Further tuning of the instrument to gentler conditions in the positive ion mode as used with the xanthine aptamer substantially increased the level of adducts observed at *m/z* regions that overlapped with the expected *m/z* of bound theophylline **4** (MW 180.16 Da) so although there may be some ligand bound it would be underneath the adduct peaks and thus difficult to discriminate, Figure 7. Both sample (RNA, ligand **4**, NH_4_OAc, DMSO, Mg^2+^, pH, refolding steps); alternate nMS buffers (trimethylammonium acetate, dimethylammonium acetate, triethylammonium acetate); alternate ions including trivalent cations; preforming the complex prior to buffer exchange; using submicrometer diameter nanospray emitters; and instrument parameter optimisation were investigated (data not shown), however there were no circumstances where theophylline **4** was observed to robustly bind to the theophylline aptamer.

**Figure 7.**
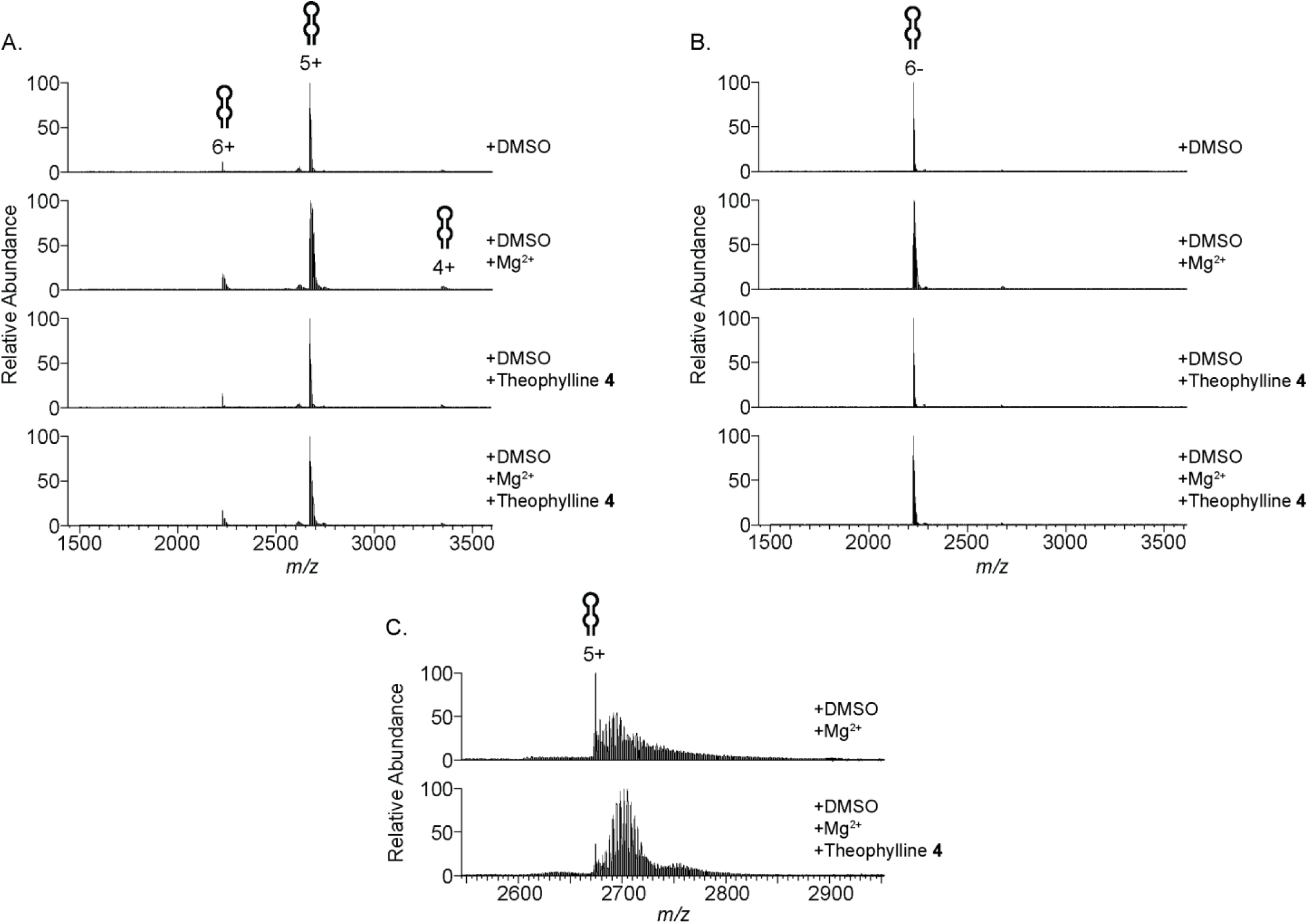
nMS analysis of theophylline RNA aptamer (2 µM, 250 mM NH_4_OAc, 1% DMSO pH 6.5), ± 20 µM Mg(OAc)_2_, ± theophylline ligand **4** in (a) positive (1 equiv) and (b) negative ion mode (1 equiv), and (c) with gentler MS instrument parameters (10 equiv). Spectrum annotations show sample conditions. Hairpin represents aptamer.

### Titration of tobramycin and kanamycin B RNA aptamers with their cognate ligands

A dose response experiment of the tobramycin and kanamycin B RNA aptamers (2 µM, 1% DMSO) with their cognate ligands, **1** and **2**, respectively (0.16-5 equiv) was performed using positive ion mode nMS with automated LC infusion (compared to static nanoESI used above). With no added ligand the tobramycin aptamer was observed as equally intense 3+ and 4+ charge states compared to predominantly the 4+ charge state observed with static nanoESI, Figure 8(a). Dose dependent binding of tobramycin **1** was observed with the measured mass shift of 467.2 Da consistent with the MW of tobramycin **1**, Table 2. The ligand binding was observed as a 4+ charged state species throughout the titration but additionally as a 3+ charge state with the higher amounts of **1** (2.5-5 equiv), Supporting Information, Figure S1. The aptamer approached saturation with 1.25 equiv of **1** and was fully saturated with 2.5 equiv of **1**, while a small amount of a second tobramycin ligand **1** binding was observed with 5 equiv of **1**. The relative intensity of 3+ charged state species (bound or unbound) decreased over the titration, possibly as the cationic aminoglycosides (positively charged at neutral pH^44^) contribute an additional positive charge to the observed species. Binding of **1** similarly occurred in the presence of Mg^2+^, Supporting Information, Figure S2(a), with the exception that at the highest titrated concentrations only the bound 4+ (and no 3+) charge state was observed (data not shown). The dimeric kanamycin B aptamer was also observed to bind to its cognate ligand kanamycin B **2** in a dose dependent manner. The measured mass shift of 483.2 Da is consistent with the MW of this ligand, Table 2. The aptamer was observed as the predominant 5+ and minor 4+ charge states and was unaltered with ligand binding, Figure 8(b) and Supporting Information, Figure S3. Ligand binding was minimal at 0.16 equiv of **2** but increased with increasing equiv of **2** with a second ligand bound at >1.25 equiv of **2** and a minor amount of a third ligand bound at 5 equiv of **2**, Figure 8(b). With addition of Mg^2+^ the extent of ligand binding remained similar, Supporting Information, Figure S2(b), however, the minor 4+ charge state was no longer visible at higher concentrations of **2** and a low intensity 6+ charge state was also present (data not shown). The nMS single point method of *K*_D_ determination^16^ was used to quantify the affinity of cognate ligand binding (as the dose responses were not sufficient to generate binding curves for *K*_D_ determination via this method), Table 4. For the first binding site of the tobramycin RNA aptamer with **1**, the *K*_D_ values across two ligand ratios (using the most abundant 4+ charge state) were 92.7 ± 16.2 nM and 74.0 ± 12.6 nM, respectively, these are similar in magnitude (only slightly weaker) than values determined via steady-state fluorescence assay data, Table 2. For the kanamycin B aptamer, *K*_D_ values across three ligand ratios (using the most abundant 5+ charge state) were consistent (786.8 ± 219.1, 845.1 ± 43.8 and 953.1 ± 126.4nM), to the best of our knowledge *K*_D_ values for cognate ligand binding to the K8-1-1kanamycin B aptamer as employed here has not previously been characterized, however, as the tobramycin RNA aptamer binding is well established this provides some confidence in the single point *K*_D_ determination method in this uncharacterized system.

**Figure 8.**
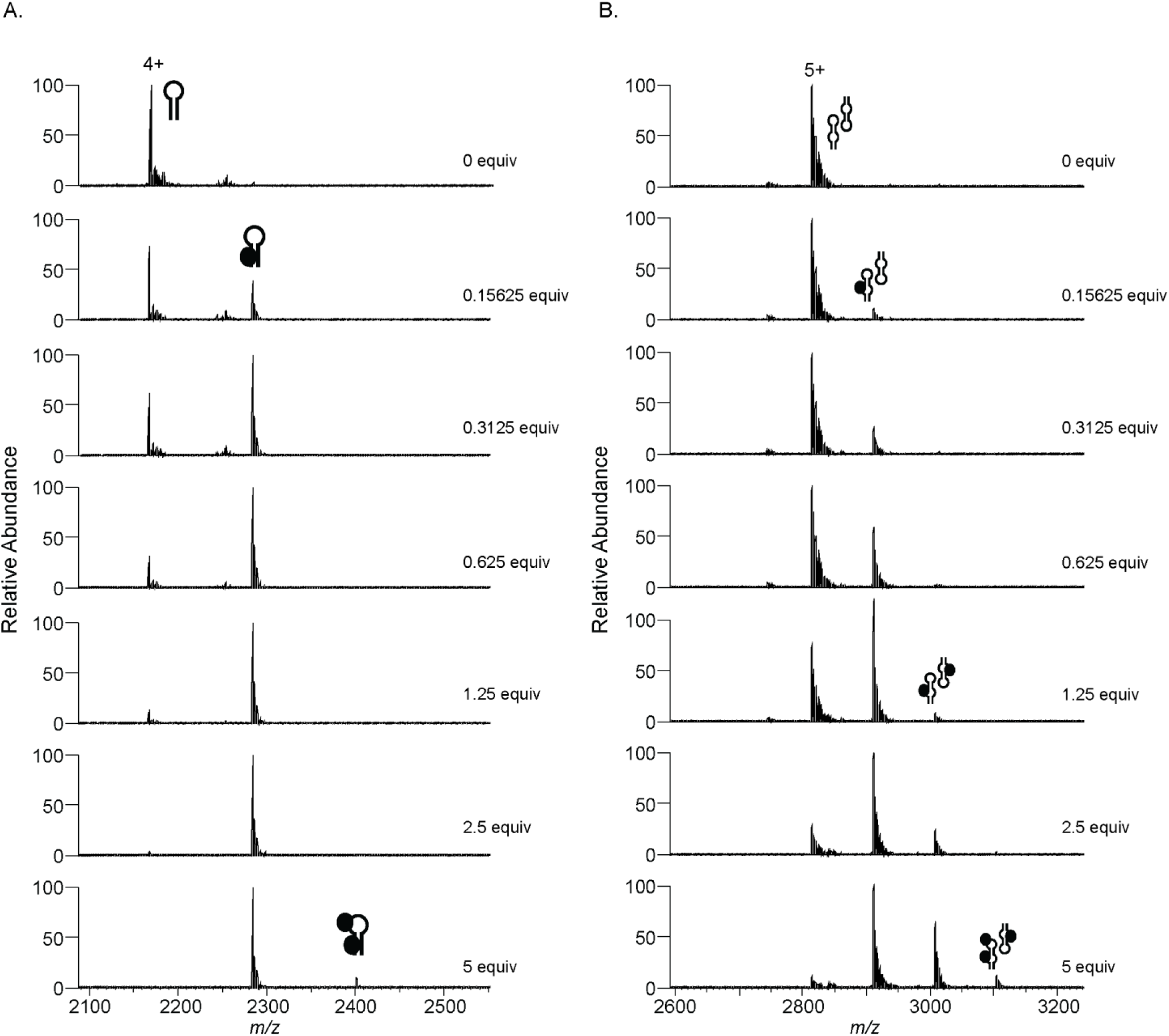
nMS analysis of dose-response binding of the cognate ligands tobramycin **1** and kanamycin B **2** to their respective aptamers (a) tobramycin RNA aptamer, and (b) kanamycin B RNA aptamer (2 µM, 250 mM NH_4_OAc, 1% DMSO pH 6.5). Spectra for the major observed charge state with ligand binding are shown. Hairpin represents aptamer, double hairpin represents dimeric aptamer, black circle represents bound ligand **1** (a) or **2** (b). nMS spectra representative of n=2.

**Table 4.**
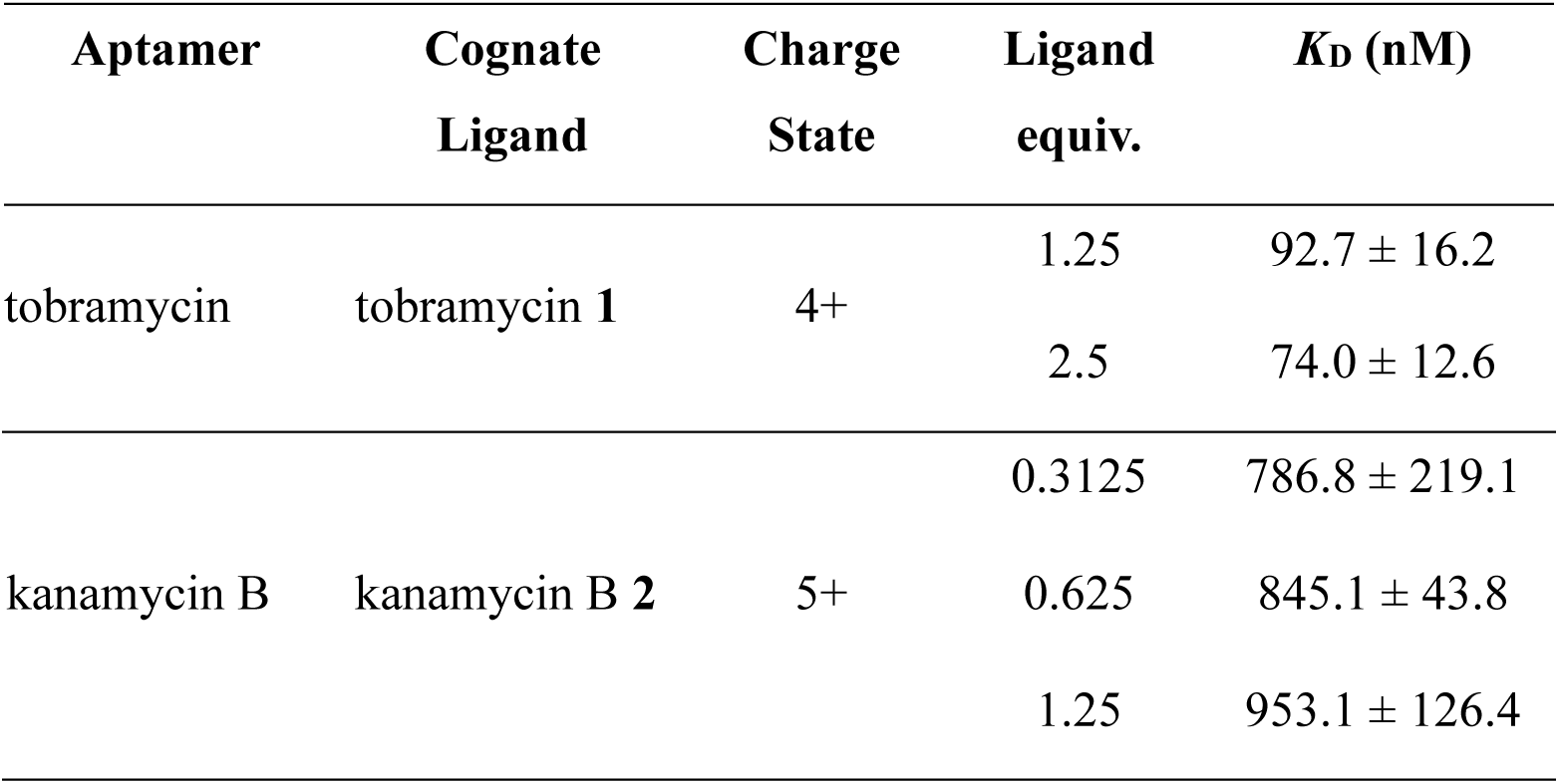
Single point *K*_D_ values (nM) as determined by nMS for the cognate ligands **1** and **2** binding their respective RNA aptamers, n=2. Single point *K*_D_ values (nM) calculated for most abundant charge state.

### Binding of tobramycin and kanamycin B RNA aptamers with the aminoglycoside panel and negative control ligands

The full aminoglycoside panel **1**, **2**, **5-8** (1 equiv) was evaluated for binding to the tobramycin and kanamycin B RNA aptamers (1% DMSO, with and without Mg^2+^) in positive ion mode. The observed charge state for binding of ligands with the tobramycin RNA aptamer was predominantly the 4+ charge state while the 4+ and 3+ charge states are of equal intensity without added ligand, with the kanamycin B aptamer ligand binding was observed as both the 5+ and 4+ charge states, with the 4+ charge state very low in intensity, Figure 9. Interestingly kanamycin B **2** was the strongest binder for both aptamers, saturating the tobramycin aptamer and over 50% bound to the kanamycin B aptamer. For the tobramycin aptamer, Kanamycin A **5** and paramomycin **7** were also strong binders (27.1 ± 0.6% and 34.2 ± 0.9% respectively), followed by neomycin **6** (13.7 ± 2.1%) and gentamicin **8** (3.4 ± 1.5%), Figure 9(a), Table 5. For the kanamycin B aptamer, paromomycin **7** and tobramycin **1** were the next strongest binders after **2** (30.3 ± 1.8% and 24.3 ± 0.6% respectively), kanamycin A **5**, neomycin **6** and gentamicin **8** were very weak binders (7.5 ± 0.1%, 9.0 ± 0.4% and 5.1 ± 3.2% respectively) Figure 9(b), Table 5. Overall, a lower level of binding of aminoglycosides was observed with the kanamycin B aptamer compared to the tobramycin aptamer, Figure 9. There was no significant difference in the aminoglycoside panel binding with Mg^2+^ present, Supporting Information, Figure S4, however, a very low intensity unbound 6+ charge state was also visible for the kanamycin B aptamer (data not shown). The nMS single point method of *K*_D_ determination^16^ was used to further quantify the affinity of aminoglycoside ligand binding, Table 5. The *K*_D_ values were calculated at 2 µM aminoglycoside ligand (using the most abundant 4+ charge state for the tobramycin aptamer, and most abundant 5+ charge state for the kanamycin B aptamer) and were consistent with the % binding, Table 5.

**Figure 9.**
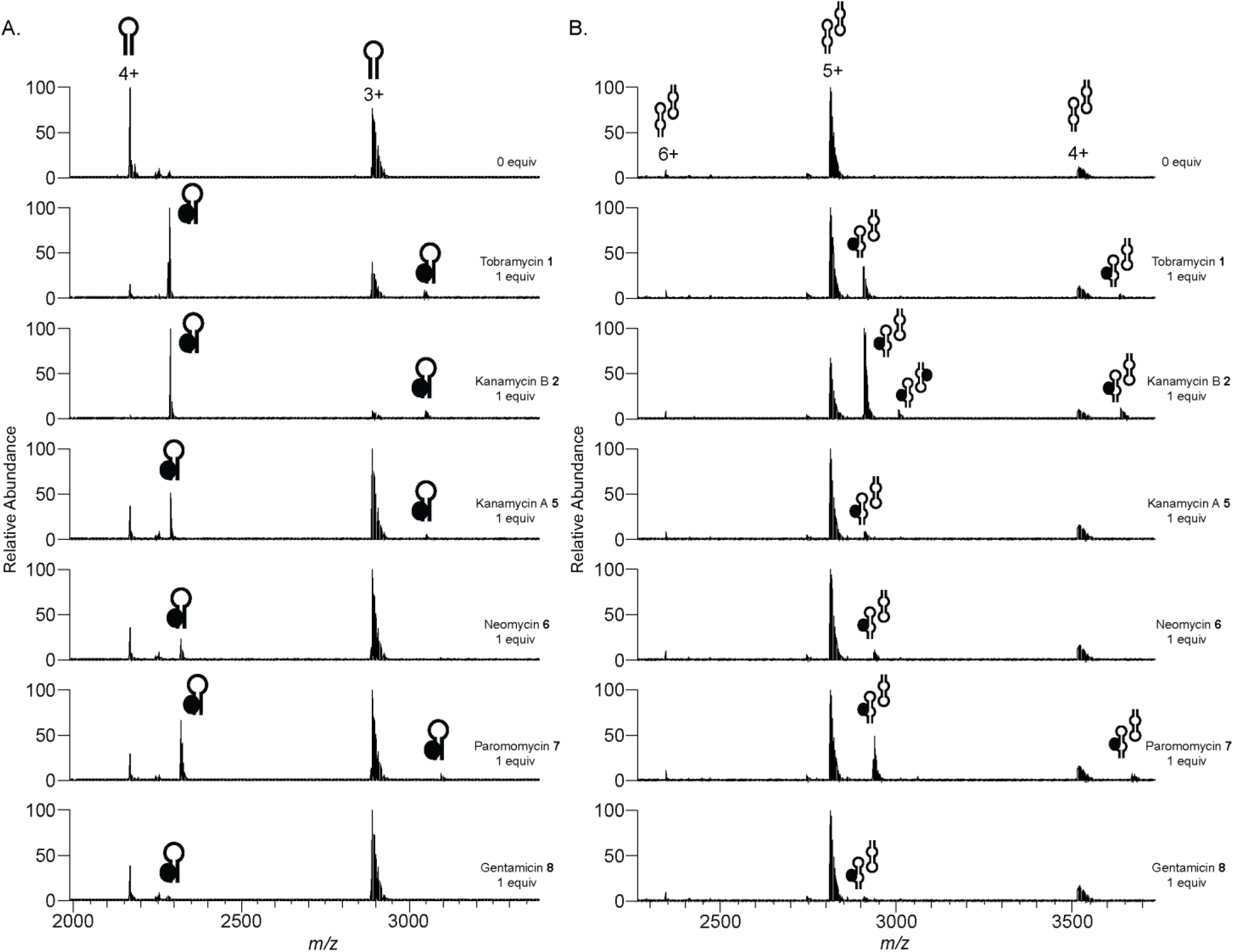
nMS analysis of the binding of the panel of aminoglycosides **1**, **2**, **5-8** to the (a) tobramycin RNA aptamer and (b) kanamycin B RNA aptamer (2 µM, 250 mM NH_4_OAc, 1% DMSO, pH 6.5) in positive ion mode with no Mg^2+^. Spectra for the major observed charge state with ligand binding are shown. Hairpin represents aptamer, black circle represents bound ligand. nMS spectra representative of n=2.

**Table 5.** Binding of the aminoglycosides **1**, **2**, **5-8** (2 µM) with the tobramycin and Kanamycin B RNA aptamers (2 µM) determined using nMS, n=2.

| | Tobramycin<br>Aptamer<br>% Binding | Kanamycin B<br>aptamer<br>% Binding | Tobramycin<br>Aptamer<br>$K_D$ (nM)<sup>a</sup> | Kanamycin B<br>aptamer<br>$K_D$ (nM)<sup>a</sup> |
| --- | --- | --- | --- | --- |
| tobramycin **1** | $66.4 \pm 0.7$ | $24.3 \pm 0.6$ | $33.7 \pm 1.0$ | $4,447.9 \pm 327.6$ |
| kanamycin B **2** | $90.1 \pm 0.7$ | $55.4 \pm 1.1$ | $2.1 \pm <0.1$ | $580.3 \pm 60.1$ |
| kanamycin A **5** | 27.1 ± 0.6 | 7.5 ± 0.1 | 603.7 ± 15.0 | 21,542.1 ± 733.5 |
| neomycin **6** | 13.7 ± 2.1 | 9.0 ± 0.4 | 1,971.4 ± 180.0 | 17,487.3 ± 683.5 |
| paromomycin **7** | 34.2 ± 0.9 | 30.3 ± 1.8 | 282.4 ± 10.9 | 2,808.2 ± 5.0 |
| gentamicin **8** | 3.4 ± 1.5 | 5.1 ± 3.2 | 13,433.95 ± 3,650.7 | 39,348.4 ± 23,434.0 |

“Single point $K_D$ values (nM) calculated for most abundant charge state (4+ for tobramycin RNA aptamer, 5+ for kanamycin B RNA aptamer).

Aptamer binding with the negative control ligand D-glucosamine **9** (1-10 equiv) as well as purine ligands caffeine **10** (10 equiv, MW 194.19 Da) and theophylline **4** (10 equiv) was assessed. A small amount of **9** was observed bound when added in excess (5-10 equiv), while no binding of caffeine **10** or theophylline **4** was observed (with or without Mg^2+^), Supporting Information, Figure S5 and S6. Collectively the findings with aminoglycoside ligands **1**, **2**, **5-8** and negative controls **4**, **9** and **10** are consistent with aptamer binding being ligand dependent, measurable using nMS and consistent with literature where available (Table 2).

### Compound library screening for binding to RNA aptamers using nMS

Significant progress has been made in curating known small molecules that bind to RNA with the development of databases that are accessible for the research community. This includes Inforna 2.0, developed by Disney and colleagues,^45^ and RNA-targeted BIoactive ligaNd Database (R-BIND and R-Bind 2.0) by Hargrove and colleagues.^46, 47^ Although the growth of both databases has been significant, with both identifying ‘RNA-privileged chemical space’, the unique small molecules within the databases are low in number (a few hundred) indicative of a need for more extensive investigations to discover RNA binding small molecules. We were interested to approach our nMS screen in an agnostic manner, specifically assessing a diverse set of small molecules for binding to the aptamers of this study to determine binders from nonbinders, relative binding strength, ligand selectivity and/or ligand promiscuity, and to assess the physicochemical and structural features of the binders as compared to nonbinders. As an agnostic approach the methods were driven by the importance for developing nMS as a robust screening method to intentionally identify RNA binders as starting points for drug discovery campaigns or for chemical probe development in academia and industry.

We curated a screening library comprising 77 compounds that were selected from the Food and Drug Administration (FDA) approved drug libraries either purchased from commercial vendors or provided by Compounds Australia (a national compound management facility supporting drug discovery),^48^ using the facility’s Smart Compounds portal^49^ to assist compound selection. Compound names, chemical structures and code numbers are in the Supporting Information, Figure S7. The screening library included 47 antibiotic drugs across seven drug classes where RNA binding is a known mechanism of action, including natural product and semisynthetic natural product drugs (seven aminoglycosides **11-17**, ten tetracyclines **18-27**, seven anthracyclines **28-34**, 12 macrolides **35-46** and two lincosamides **47-48**) and synthetic drugs (six fluoroquinolines **49-54** and three oxazolodinones **55-57**).^24^ Additionally 30 drugs without a RNA binding pedigree were selected to accommodate compounds with physicochemical parameters that predominantly fall within Lipinksi’s rule of five guidelines^27^ or fragment guidelines,^50^ these comprised two subsets – 15 drugs of MW >300 Da **58-72** and 15 fragment sized drugs of MW <300 Da **73-87**. As the chemical and structural characteristics of RNA differ markedly to proteins, these lower complexity drug-like compounds were of interest because they are in principle more readily developable by medicinal chemistry programs. Selected physicochemical properties for the screening library including MW, calculated Log P (clog P), topological polar surface area (PSA), number of H-bond acceptors, number H-bond donors, number of chiral centres and number of rotatable bonds are in Table 6, Figure 10. The wide distribution of properties highlights diversity in the screening library compounds.

**Table 6.**
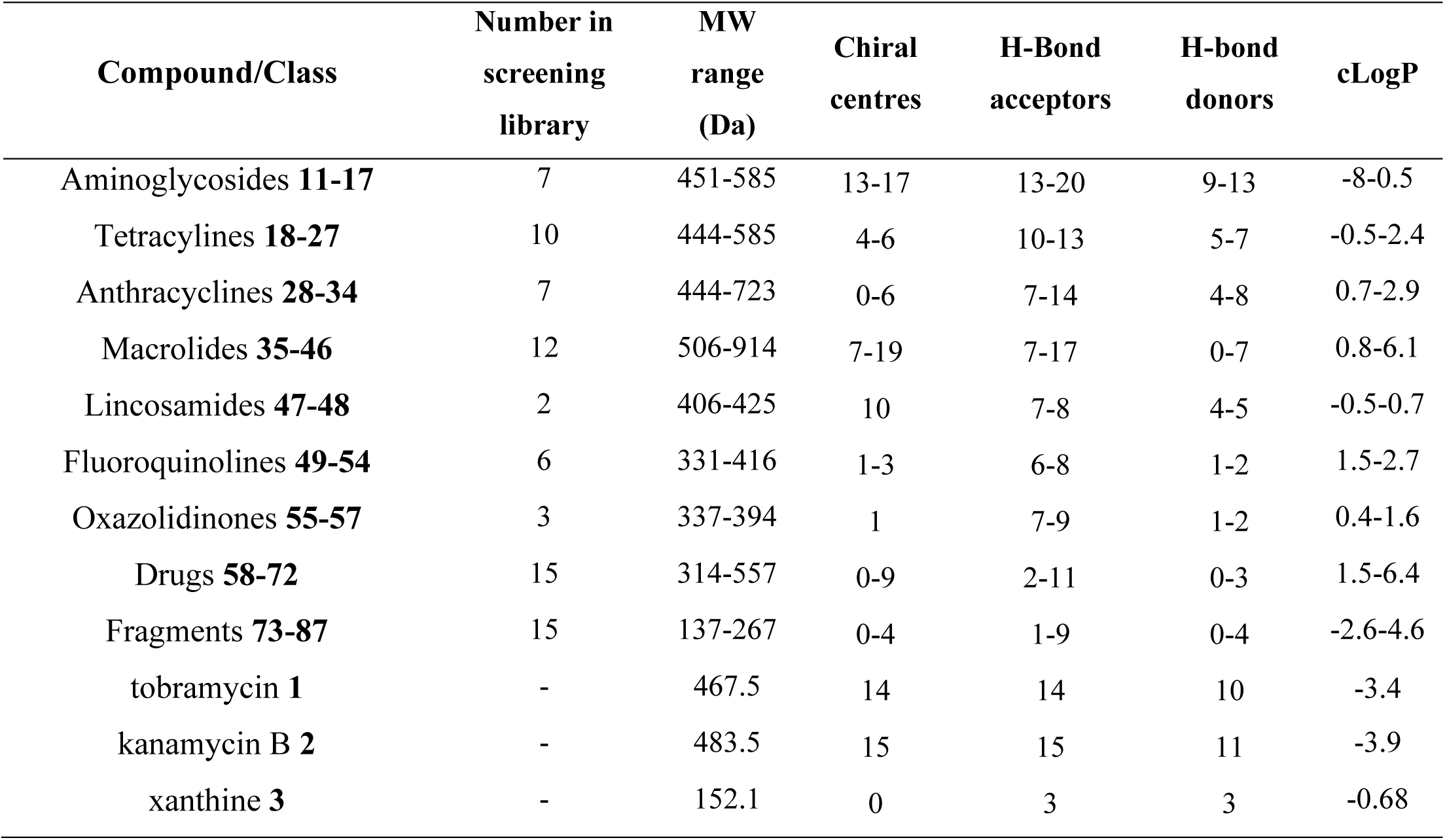
Range of selected physicochemical properties of the 77-member drug screening compound library **11-87** and aptamer cognate ligands **1-3** used for screening against RNA aptamers using nMS.

**Figure 10.**
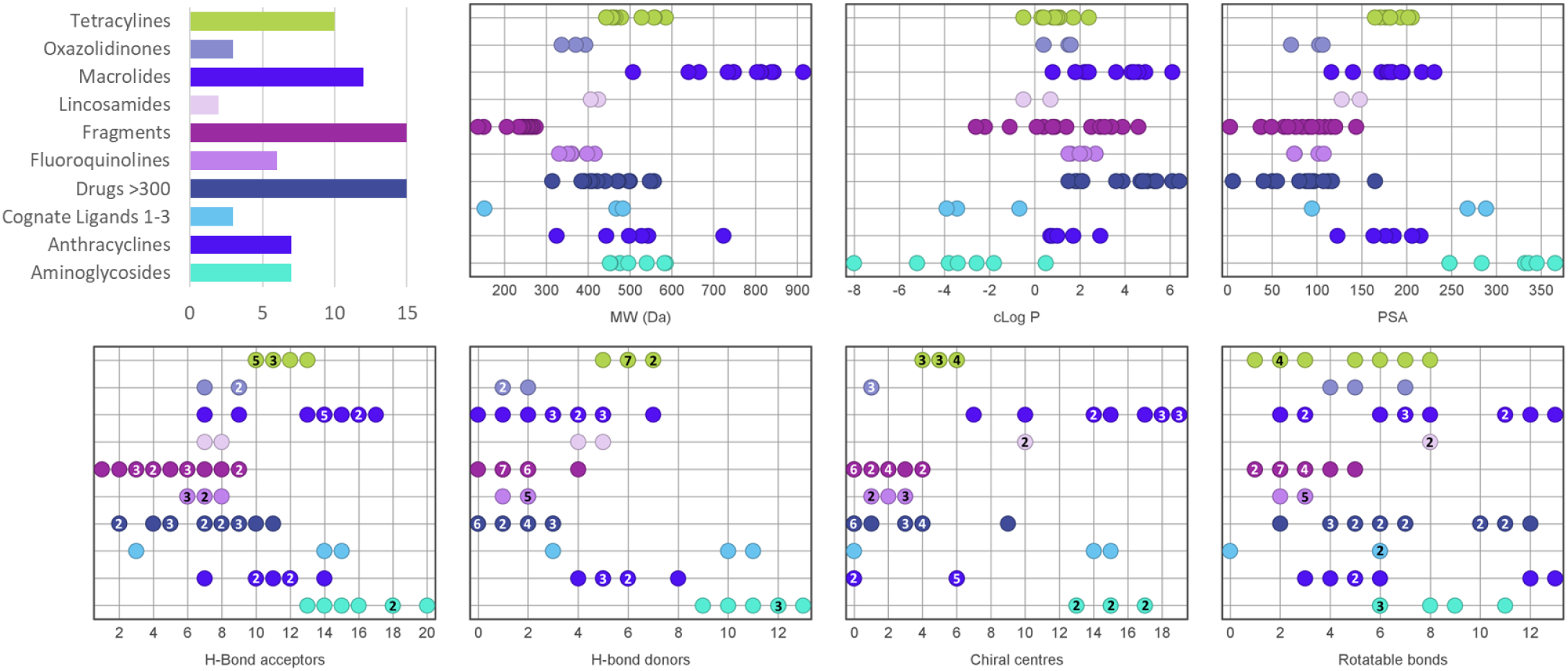
Distribution of selected physicochemical properties for the nine compound classes within the drug screening library (compounds **11-87**). Properties include MW, and calculated Log P (clog P), topological polar surface area (PSA), number of H-bond acceptors, number H-bond donors, number of chiral centres and number of rotatable bonds. Where there is more than one compound with the same value this is indicated by inclusion of a number within the circle. Properties for the cognate aptamer ligands **1-3** is included for comparison. Figure generated using DataWarrier.^51^

The compound library (5 µM, 2.5 equiv) was screened using nMS in positive ion mode against the tobramycin, kanamycin B and xanthine RNA aptamers (2 µM). Tobramycin **1**, kanamycin B **2** and xanthine **3**, Figure 2, were included in the screening protocol as positive controls (2.5 equiv) for their corresponding aptamer. As Mg^2+^ was not required for cognate ligand or aminoglycoside panel binding studies described above, the RNA aptamer screening was performed without added Mg^2+^. Samples were incubated for a minimum of 30 min at room temperature prior to nMS analysis to enable equilibration of the RNA-ligand noncovalent complexes. nMS data was deconvoluted using UniDec^52^ and ligand binding quantified using a Python script as reported by us previously.^53^ In brief, the script identifies and normalizes the intensity of the most intense peaks from the deconvolution to sum to 100%, enabling determination of % ligand binding, Table 7, and producing a heat map to easily visualize deconvoluted spectra and identify binding, Figure 11. The chemical structures of the screening hits are in Figure 12. Tobramycin **1** saturated the tobramycin aptamer with a small amount of aptamer observed with two ligands bound (2.6%); kanamycin B **2** bound to the dimeric aptamer (60.1% with one ligand bound and 14.5% with two ligands bound), while binding of xanthine **3** to the xanthine aptamer was not observed with the screening conditions employed (screening MS parameters were not the gentler parameters required to observe xanthine binding as described above but were the parameters to decrease adducts and improve screening sensitivity). All aminoglycosides **11-17** in the screening library bound to the tobramycin and kanamycin B aptamers, however, none bound more strongly than the cognate aminoglycoside ligands **1** or **2**. Binding was observed with either one or two ligands binding to the tobramycin aptamer and one to four ligands observed binding to the dimeric kanamycin B aptamer. Geneticin **14** was the weakest binder (11.9% and 8.0%, respectively) and all aminoglycosides except for geneticin also had measurable binding to the xanthine aptamer. Aminoglycosides dibekacin **13** and netilmicin **15** were the strongest xanthine aptamer binders and exhibited a high amount of multiple binding across all three RNA aptamers, which indicates poorly selective ligands and/or nonspecific binding, this is consistent with the reported promiscuous binding of aminoglycosides to RNA^25^ and with other aminoglycoside compounds **5-8** as described above. However, the % binding data shows that there is a measurable level of discrimination in binding within the class and between the different RNA aptamers, as well as confirms stronger binding of aminoglycosides compared to other compound classes, Table 5, collectively reinforcing the importance of methods that can measure and quantify RNA binding and binding differences. The core structure of aminoglycosides consists of a 2-deoxystreptamine or streptamine moiety with amino sugar substituents at different positions.^24^ The mechanism of action is driven by the presentation of basic amino groups and hydrogen bond donor moieties that participate to provide a scaffold for a network of interactions with the ribosomal RNA targets,^23^ while it has been demonstrated that both electrostatics and conformational flexibility play a significant role in the ability of aminoglycosides to bind to RNAs more generally.^4^ Further, as development of RNA aptamers provides many RNA sequences that can bind the cognate ligand (e.g. in the vicinity of 10^7^-10^8^ RNA sequences in the selected pool that could bind tobramycin **1**^31^) this implies that an aminoglycoside ligand can bind to a large number of alternative RNA sequences with moderately high affinity.

**Table 7.** Binding of control compounds and hits (5 µM, 2.5 equiv) with the tobramycin, Kanamycin B and xanthine RNA aptamers (2 µM) determined using nMS.

| Class | Drug | MW | Average % binding |  |  |  |  |  |  |  |  |
| --- | --- | --- | --- | --- | --- | --- | --- | --- | --- | --- | --- |
|  |  |  | Tobramycin Aptamer |  | Kanamycin B Aptamer |  |  |  | Xanthine Aptamer |  |  |
|  |  |  | +1 Ligand | +2 Ligands | +1 Ligand | +2 Ligands | +3 Ligands | +4 Ligands | +1 Ligand | +2 Ligands | +3 Ligands |
| Cognate ligands | Tobramycin **1** | 467.50 | 97.4 ± 0.7 | 2.6 ± 0.7 | | | | | | | |
| Cognate ligands | Kanamycin B **2** | 483.50 | | | 60.1 ± 0.1 | 14.5 ± 0.1 | | | | | |
| Cognate ligands | Xanthine **3** | 152.10 |  |  |  |  |  |  | * |  |  |
| Aminoglycosides | Amikacin **11** | 585.61 | 59.9 ± 2.8 | 6.9 ± 4.9 | 32.3 ± 1.0 | 7.2 ± 1.6 | | | 12.6 ± 0.6 | | |
| Aminoglycosides | Apramycin **12** | 539.58 | 64.0 ± 12.2 | 31.5 ± 18.7 | 43.4 ± 0.4 | 32.0 ± 0.5 | 8.9 ± 0.4 | | 22.6 ± 2.5 | | |
| Aminoglycosides | Dibekacin **13** | 451.52 | 33.0 ± 2.0 | 64.8 ± 1.2 | 23.9 ± 6.2 | 38.7 ± 2.8 | 28.0 ± 5.0 | 6.1 ± 3.5 | 34.1 ± 0.1 | 34.8 ± 1.1 | 5.8 ± 0.6 |
| Aminoglycosides | Geneticin **14** | 496.55 | 11.9 ± 0.0 | | 8.0 ± 0.6 | | | | | | |
| Aminoglycosides | Netilmicin **15** | 475.59 | 54.4 ± 9.1 | 41.9 ± 15.6 | 16.1 ± 3.0 | 26.8 ± 2.8 | 28.5 ± 0.8 | | 24.5 ± 0.8 | 37.2 ± 0.8 | 12.3 ± 1.1 |
| Aminoglycosides | Ribostamycin **16** | 454.48 | 55.9 ± 12.4 | 6.2 ± 5.4 | 24.6 ± 1.1 | 3.2 ± 0.4 | | | 10.6 ± 1.2 | | |
| Aminoglycosides | Streptomycin **17** | 581.58 | 27.2 ± 3.8 | 16.5 ± 11.0 | 8.7 ± 1.4 | 25.3 ± 5.3 | | | 7.3 ± 3.5 | 5.0 ± 3.3 | |
| Anthracyclines | Pixantrone **28** | 325.37 | 60.1 ± 14.3 | | | | | | 13.0 ± 8.6 | | |
| Anthracyclines | Daunorubicin **29** | 527.53 | 19.6 ± 1.8 | | 5.9 ± 1.8 | | | | 4.3 ± 0.1 | | |
| Anthracyclines | Doxorubicin **30** | 543.53 | 17.2 ± 0.4 | | 5.1 ± 2.8 | | | | 4.4 ± 0.0 | | |
| Anthracyclines | Epirubicin **31** | 543.53 | 19.3 ± 0.1 | | 4.5 ± 1.3 | | | | 5.0 ± 0.4 | | |
| Anthracyclines | Idarubicin **32** | 497.50 | 4.3 ± 1.3 | | | | | | 1.5 ± 0.1 | | |
| Anthracyclines | Mitoxantrone **33** | 444.20 | 52.8 ± 15.5 | | 19.7 ± 8.1 | | | | 24.8 ± 6.4 | | |
| Macrolides | Ixabepilone **40** | 506.70 | 9.0 ± 7.5 | | | | | | | | |
| Macrolides | Tacrolimus **43** | 804.03 | 9.4 ± 3.3 | | | | | | | | |
| Macrolides | Rapamycin **39** | 914.19 | 2.0 ± 0.8 |  |  |  |  |  |  |  |  |
| Oxazolidinones | Eperezolid **55** | 394.40 | 8.6 ± 1.2 |  | 2.7 ± 0.6 |  |  |  |  |  |  |
| Drugs >300 | Flumethasone **60** | 410.46 | 17.8 ± 2.1 |  | 5.6 ± 1.6 |  |  |  | 2.0 ± 0.6 |  |  |
| Drugs >300 | Eszopiclone **62** | 388.81 | 9.1 ± 3.7 |  | 3.6 ± 1.8 |  |  |  |  |  |  |
| Drugs >300 | Edoxaban **68** | 548.06 | 14.3 ± 0.8 |  | 3.9 ± 2.0 |  |  |  |  |  |  |
| Drugs >300 | Linagliptin **70** | 472.55 | 11.0 ± 0.7 |  | 1.4 ± 2.0 |  |  |  |  |  |  |
| Drugs >300 | Rolapitant **72** | 500.49 | 11.9 ± 3.1 |  | 3.8 ± 0.4 |  |  |  | 3.2 ± 0.3 |  |  |
| Fragments | Dapsone **76** | 248.30 | 5.6 ± 4.2 |  |  |  |  |  |  |  |  |

Data is average ± standard deviation, n=2. Dark grey shading denotes compound not tested against this aptamer. Light grey shading denotes binding not observed.
\*no binding of the cognate ligand **3** to the xanthine aptamer at 2.5 equiv with the nMS parameters employed for screening.

**Figure 11.**
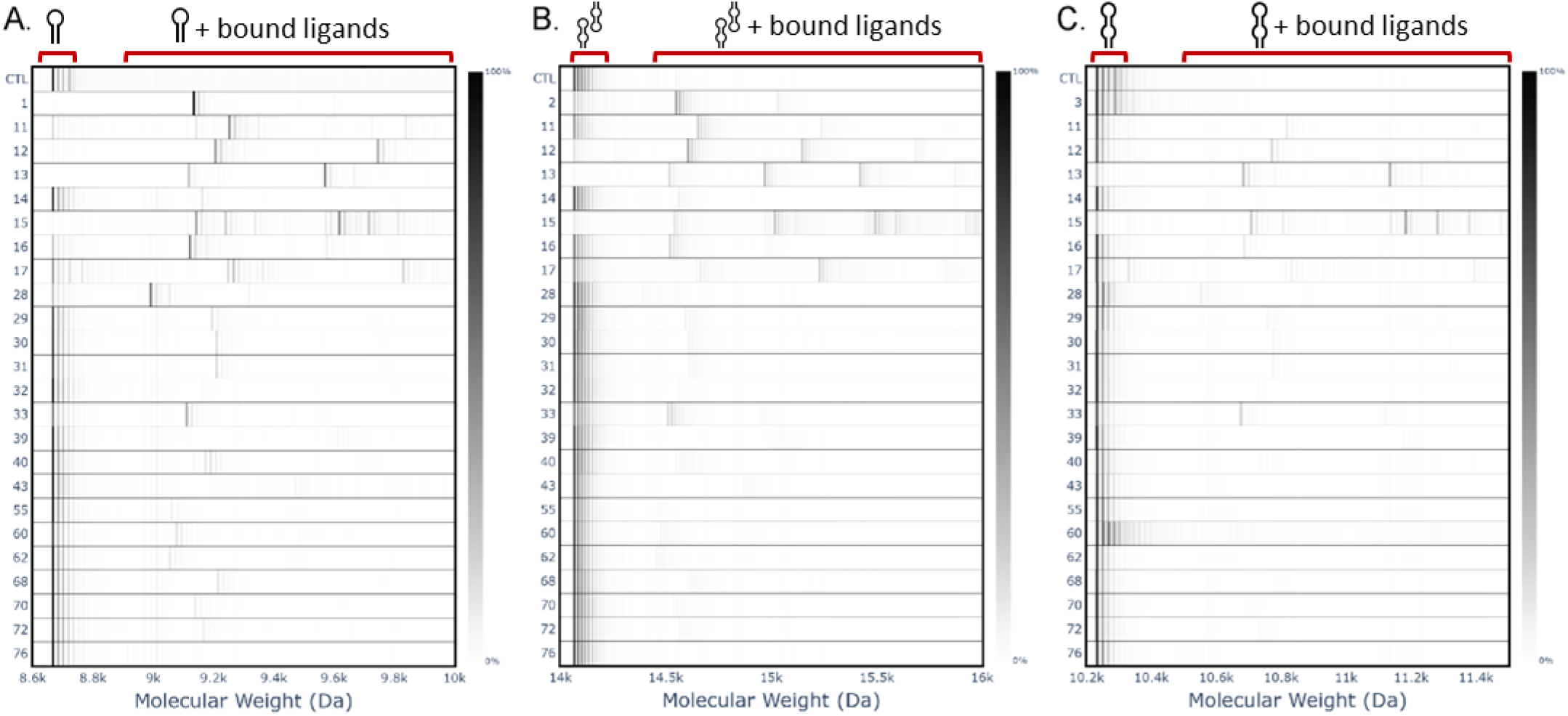
nMS heat maps of the deconvoluted native mass spectra for screening hits acquired from screening RNA aptamers against library **11-87**. The aptamer plus bound ligand peak intensity is indicative of ligand binding strength with quantitative data in Table 5. (a) With tobramycin RNA aptamer. (b) With kanamycin B RNA aptamer. (c) With xanthine RNA aptamer. N=2 (1 replicate only shown). CTL is the RNA aptamer only control sample. Cognate ligands **1-3** are positive controls for the corresponding RNA aptamer. Screening hit ligand code number are on y axis.

**Figure 12.**
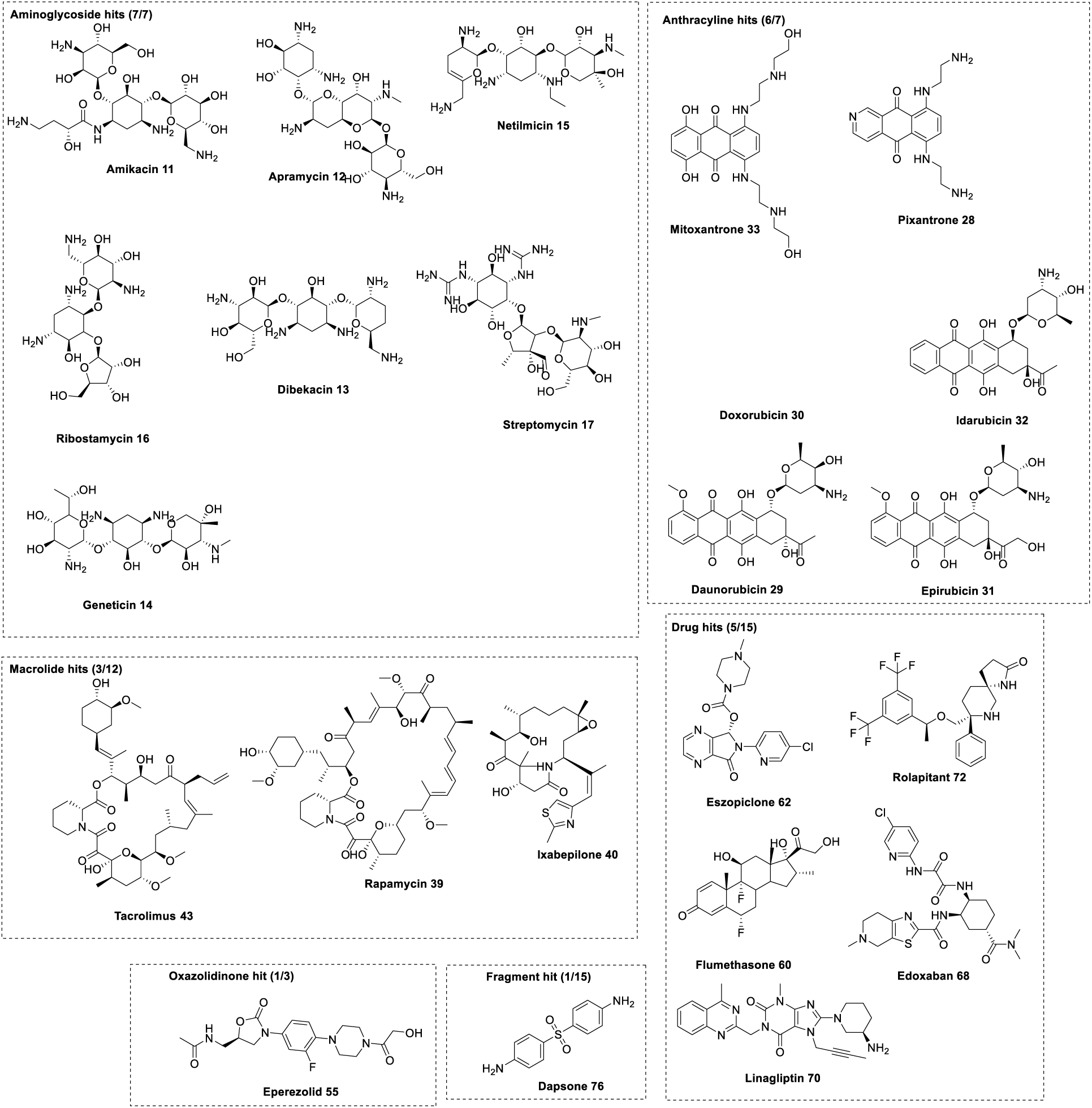
Screening hits of library of FDA approved drugs to RNA aptamers using nMS.

Anthracyclines were the next most dominant class of RNA aptamer binders after aminoglycosides, with six of the seven anthracyclines binding across the three aptamers but with stronger binding to the tobramycin aptamer. Mitoxantrone **33** and pixantrone **28**, closely related structures, were the strongest binders (52.8% and 60.1% respectively to the tobramycin aptamer) and were similar in binding strength as the aminoglycosides. Other anthracyclines were moderate binders (∼20% bound to the tobramycin aptamer) with one (idarubicin **32**) weakly bound (∼4% bound to the tobramycin aptamer). Three macrolides (ixabelipone **40**, tacrolimus **43** and rapamycin **39**) bound weakly and selectively to the tobramycin aptamer (∼2-9% binding) with the remaining nine macrolide representatives nonbinders. One of three oxazolidinones bound weakly and selectively to the tobramycin aptamer (eperozolid **55**, ∼8% binding). Of the fragment class only one compound, dapsone **76** – a bis aniline sulfoxide with no chiral centres – had weak but selective binding to the tobramycin aptamer (∼6% binding) and may be interesting to further study if this may be a privileged scaffold for RNA binding and inclusion in RNA screening libraries. Five of the compounds in the drug class (>300 Da) exhibited binding across the different aptamers with the trend of stronger binding to the tobramycin aptamer observed as with the other compound classes. Of these, two of the drugs (rolapitant **72** and flumethasone **60**) displayed some binding to all aptamers, while three (eszopiclone **62**, edoxaban **68** and linagliptin **70**) bound only to the tobramycin and kanamycin B aptamers. The structures of the hit drugs as well as nonbinding drugs are diverse, and the binding and selectivity for some of this group is promising and supports the benefit of wider consideration of FDA approved small molecule drugs in screening libraries against RNA drug targets providing insightful data for potential off target effects or new leads for development as chemical probes, repurposed drugs or novel new drugs. Finally, binding of the tetracyline (10 examples), fluoroquinoline (6 examples) and lincosamide (2 examples) compound classes was not observed with any of the RNA aptamers, this is also important as it affirms discrimination of small molecule binding to RNA, and that this is readily measurable with nMS.

The physicochemical properties for the screening hits are provided in Supporting Information, Figure S8, with properties of stronger hits (>20 % binding) summarized here as they may be indicative of those with favoured RNA binding. Stronger hit properties include MW all >300 Da but <600 Da, number of hydrogen bond donors all >4, number of hydrogen bond acceptors all >6, topological polar surface area (PSA) all >100 and calculated Log P (clog P) all <2. The number of chiral centres was, however, 0 to 17 (indicative of potential intercalators when 0 and more complex shape/electrostatic complementarity interactions when high in number). These physicochemical properties fit with the average values where reported by Disney and colleagues.^8^ Furthermore, aminoglycosides are the predominant class with >20% binding, and when compared with FDA approved drugs more generally, have lower clog P values, higher PSA values, and high hydrogen bond acceptor and hydrogen bond donor count.^8^

## CONCLUSION

nMS has proven a powerful biophysical technique to characterize protein-ligand binding for primary screening of small molecules. Herein we have explored the use of nMS to characterize the binding of small molecules with model RNA aptamers, demonstrating that nMS may be expanded to native RNA targets for small molecule library screening, where robust and orthogonal methods are needed to support new therapeutic development. The nMS workflow is target agnostic and importantly provides information on ligand binding, strength of binding, specificity and selectivity towards RNA, while other benefits include low sample amount requirements, scalable to high throughput and direct detection without modifications of the RNA. The nMS platform was applicable in both positive ion mode and negative ion mode across a range of sample and instrument parameters, enabling a flexible workflow for both single point and dose response binding measurements. This work validated the limited reported findings of the quantitative and qualitative binding with the aminoglycoside class of antibiotics. By comparative screening of three different RNA aptamers as targets, we found that ligands, inclusive of a library of FDA approved drugs (with and without known RNA binding pedigrees) were detectable with nMS with potential chemical property preferences for binding and nonbinding ligands identifiable, while nonselective ligands were readily distinguished from more specific ligands. A key challenge of the nMS method with RNA is the high level of non-structural salt adducts that can lower the sensitivity of detection of ligand binding and can hinder accurate binding quantification, however, the RNA ligand bound nMS peaks were distinct from the RNA adduct peaks with the instrument parameters tuned for screening. As the nMS of high salt samples can be improved using nanoelectrospray emitters with submicron diameter openings, exploring this approach in future studies could be of interest.^42^ In summary, the RNA aptamers selected for this study displayed expected binding affinity for their cognate ligands, with specificity of binding to nine diverse ligand classes measurable, spanning strong, weak or no binding. It is expected that nMS will enable efficient, target-based yet target-agnostic characterization of both specific and off-target RNA–small molecule interactions supporting future screening campaigns, providing evidence of target engagement and an alternative approach to generate or validate structure-activity relationships.

## EXPERIMENTAL

### RNA aptamers and test compounds

Synthetic RNA aptamers were purchased from BioSpring (theophylline, kanamycin B and tobramycin RNA aptamers) or IDT (xanthine RNA aptamer) and stored in lyophilized form at 4 °C. Stock solutions of the RNA aptamers 100-200 µM were prepared in nuclease-free distilled deionized water, aliquoted and stored at 4 °C. The nMS buffer components including nuclease-free water and nuclease-free ammonium acetate (5 M solution) were purchased from Thermo Fisher Scientific, and DMSO from Sigma Aldrich. Control small molecule ligands were purchased from AK Scientific or Sigma Aldrich (Supporting Information, Table S1). Stock solutions of ligands 1-10 were prepared in either 50% or 100% DMSO or H_2_O (5-20 mM), depending on the solubility of the compound, and stored at -20 °C. Compounds were diluted to working stocks that were 10-fold more concentrated than the required screening concentrations with final 10% DMSO. The small molecules for screening against the RNA aptamers were supplied by Compounds Australia, Griffith University or purchased from AK Scientific or Sigma Aldrich (Table S1). Specifically, 200 nL of 5 mM compound stocks in 100% DMSO were plated in nMS-compatible 96-well plates (BioRad Hard-Shell® 96-well PCR plates #HSP9601). Column 1 was plated with 200 nL DMSO as a negative control, and column 2 plated with 200 nL of the cognate ligand as a positive control. This plate layout provided of a positive and negative control for every ten test compounds and thus allowed monitoring of the stability of the nMS screening assay. Additional details of the compounds 11-76 selected for screening against the RNA aptamers are provided in the Supplementary Information.

### Native mass spectrometry sample preparation

Prior to nMS analysis, an aliquot of the RNA aptamer stock was buffer exchanged into 250 mM NH_4_OAc, pH 6.5 using Amicon Ultra-0.5 3 kDa cut-off centrifugation units. Following buffer exchange the concentration of the RNA aptamer sample was determined by measuring the absorption at 260 nm using a NanoDrop microvolume UV-Vis spectrophotometer (Thermo Fisher Scientific, Bremen, Germany) and adjusted to 2 µM by addition of 250 mM NH_4_OAc, pH 6.5. A native mass spectrum of each aptamer (2 µM, 250 mM NH_4_OAc, ±20 µM Mg(OAc)_2_, ± 1% (v/v) DMSO, pH 6.5) was acquired in both the positive ion and negative ion mode with and without 1 equiv of the cognate ligand. For the ligand binding studies, 2 µM RNA aptamer (18 µL) was incubated with 1 equiv of the ligand (2 µL) at room temperature for at least 30 min prior to nMS analyses. The 6-point dose-response experiments were conducted using 2 µM RNA aptamer (18 µL) with the addition of 2 µL of the cognate ligand for each aptamer at RNA to final ligand concentration ratios of 0.16-5 equiv. The negative control compounds D-glucosamine 9, theophylline 4 and caffeine 10 were each tested against 2 µM RNA at 1, 5 and 10 equiv. Samples were incubated at room temperature for at least 30 min prior to nMS analyses.

For compound screening the corresponding RNA aptamer (2 µM) was added to each well of the multiwell plate containing the screening compounds (described above) to yield samples with 2.5 equiv, 5 µM ligand. All samples were incubated for 30 min at room temperature prior to nMS analysis.

### Native mass spectrometry instrumentation

The cognate ligand studies for all aptamers were acquired using a Q-Exactive UHMR mass spectrometer (Thermo Fisher Scientific) fitted with a Nanospray Flex ion source with static spray head. Data was collected in both positive and negative ion mode using nanoESI from platinum-coated borosilicate capillaries purchased from Thermo Fisher Scientific (inner diameter ∼10-15 µm, ES388). Positive ion nMS parameters were as follows: spray voltage 1.1 kV (for tobramycin and kanamycin B aptamers), 1.5 kV (for xanthine aptamer) and 0.9 kV (for theophylline aptamer), resolution 200,000, microscans 5, no averaging, S-lens RF level at 200, extended trapping 1.0 eV, desolvation voltage 0 V (for tobramycin aptamer) and –15 V (for kanamycin B, xanthine and theophylline aptamers), trapping gas pressure setting 1.0, capillary temperature 275 °C, detector *m/z* optimisation set for low *m/z*, ion transfer target *m/z* set for high *m/z*, scan range 1,500-6,000 *m/z*, data acquisition length for at least 0.5-2 min. Negative ion nMS parameters were: spray voltage 1.9 kV (for tobramycin aptamer), 2.1 kV (for kanamycin B and xanthine aptamers) and 1.3 kV (for theophylline aptamer), resolution 200,000, microscans 5, no averaging, S-lens RF level at 200, extended trapping 1.0 eV, desolvation voltage 0 V (for tobramycin aptamer) and –15 V (for kanamycin B, xanthine and theophylline aptamers), trapping gas pressure setting 1.0, capillary temperature 275 °C, detector *m/z* optimisation set for low *m/z*, ion transfer target *m/z* set for high *m/z*, scan range 1,500-6,000 *m/z*, data acquisition length for at least 0.5-2 minutes.

Compound screening was performed using a Vanquish Neo UHPLC system (Thermo Fisher Scientific) coupled to a Q-Exactive UHMR mass spectrometer (Thermo Fisher Scientific) to acquire native mass spectra. The micro-flow LC system conditions were 250 mM NH_4_OAc pH ∼6.5 used as the solvent mobile phase for direct sample injection of 2 µL from each assay plate well at 10 µL/min. The sample was infused into the mass spectrometer using the Nanospray Flex ion source fitted with the direct junction and via a stainless steel nanobore emitter (internal diameter 30 µm, ES542). MS parameters were as follows: positive polarity, spray voltage 1.8 kV, resolution 200,000, microscans 5, no averaging, S-lens RF level at 200, extended trapping 1.0 eV, desolvation voltage 0 V (for tobramycin aptamer) and –15 V (for kanamycin B and xanthine aptamers), trapping gas pressure setting 1.0, capillary temperature 275 °C, ion detection optimised for low *m/z*, ion transmission optimised for high *m/z*, scan range 1,500-6,000 *m/z*. Data were acquired for 2 min for each sample (allowing sample injection, acquisition of spectrum and the washing away of sample with solvent) with sample carryover minimal.

### Native mass spectrometry data analysis

#### Aptamer only

Data analyses were completed manually in FreeStyle (Thermo Fisher), with aptamer MW calculated from the *m/z* of the unadducted peak of the most intense charge state using the following equation MW = (*m/z*-1) × *z*.

#### Aptamer ligand binding and screening

Mass spectrometry screening data was analysed by batch deconvolution in UniDec^52^ followed by an inhouse python script to visualize the deconvoluted MS spectrum as a heatmap enabling ease of identification of ligand binding (or hits), as previously reported.^53^ Data analysis parameters for each aptamer were as follows: (a) tobramycin RNA aptamer UniDec parameters: 1,700-3,500 *m/z* range, 3-6 charge state range, 8,500-10,000 Da MW range, sample mass every 1.0 Da, peak detection range and threshold of 100 Da and 0.1 respectively; script parameters: 8,500-10,000 Da MW range; (b) kanamycin B RNA aptamer UniDec parameters: 2,000-4,000 *m/z* range, 3-6 charge state range, 14,000-16,000 Da MW range, sample mass every 1.0 Da, peak detection range and threshold of 100 Da and 0.1 respectively; script parameters: 14,00-16,000 Da MW range, (c) xanthine RNA aptamer UniDec parameters: 2,200-3,700 *m/z* range, 3-5 charge state range, 10,200-12,000 Da MW range, sample mass every 1.0 Da, peak detection range and threshold of 100 Da and 0.1 respectively; script parameters: 10,200-11,500 Da MW range.

As nMS resolves unbound RNA from ligand bound RNA, quantification of the relative abundance of ligand binding to the RNA aptamer was determined from the UniDec^52^ deconvoluted spectra by the python script as previously described.^53^ Briefly, the five most intense peaks in the native mass spectrum following UniDec^52^ deconvolution were normalized to sum to 100%. The script labels these deconvoluted peaks with their molecular weight allowing the molecular weight of the unbound RNA and ligand bound RNA (when present) to be determined and, thus, also ligand identity to be confirmed. Additionally, the script provides the percentage of ligand binding relative to unbound RNA. Single point *K*_D_ values were also determined for the binding of control ligands according to previously published methods/equations,^16^ and using the absolute intensity values obtained from the raw mass spectra for the most abundant charge states where ligand binding was observed.

## ACKNOWLEDGEMENTS

The authors gratefully acknowledge support of the Ramaciotti Foundations (Biomedical Research Award, grant number 2023BRA19) and the Australian Government through an Australian Research Council Industrial Transformation Training Centre grant (grant number IC180100021), and Australian Research Council Linkage, Infrastructure, Equipment and Facilities grants (grant numbers LE120100170, LE220100031). We acknowledge Compounds Australia (www.compoundsaustralia.com) for their provision of specialized compound management and logistics research services to the project. We acknowledge Mike Baksh, Sean Harrison, Lilianna Pedro, Hamed Pirimoghadam, Elisa Barile, Pedro Serrano of Takeda for helpful discussions and providing theophylline, kanamycin B and tobramycin RNA aptamer samples. We thank Yezhou (Frank) Yu for support with application of the python script. We thank Dr Bruno Madio for assistance with instrumentation installation and setup. We thank A/Prof Michael Marty and James Sanders, Prof W. Alexander Donald and Manatsu Nose for helpful discussions on nMS and Dr Maria Halili for discussions on RNA samples and handling, and experimental/project set up.

## SUPPORTING INFORMATION

Sample suppliers of compounds; nMS spectra for aptamers and ligands; physicochemical properties of hits and chemical structures for full compound library.

## AUTHOR CONTRIBUTIONS

SAP and LS conceived this study and methodology. LS and JK performed the experiments. SAP supervised the project. All authors analyzed and curated the data. All authors contributed to the preparation of the manuscript. SAP was responsible for funding acquisition for the project.

## Notes

### Competing Interest Statement

The authors have declared no competing interest.

## REFERENCES

(1) Santos, R.; Ursu, O.; Gaulton, A.; Bento, A. P.; Donadi, R. S.; Bologa, C. G.; Karlsson, A.; Al-Lazikani, B.; Hersey, A.; Oprea, T. I.;, et al. A comprehensive map of molecular drug targets. Nat Rev Drug Discov 2017, 16 (1), 19–34. DOI: 10.1038/nrd.2016.230 From NLM.

(2) Warner, K. D.; Hajdin, C. E.; Weeks, K. M. Principles for targeting RNA with drug-like small molecules. Nat Rev Drug Discov 2018, 17 (8), 547–558. DOI: 10.1038/nrd.2018.93.

(3) Haniff, H. S.; Knerr, L.; Chen, J. L.; Disney, M. D.; Lightfoot, H. L. Target-Directed Approaches for Screening Small Molecules against RNA Targets. SLAS Discovery 2020, 25 (8), 869–894. DOI: 10.1177/2472555220922802 (acccessed 2024/08/29).

(4) Ratni, H.; Scalco, R. S.; Stephan, A. H. Risdiplam, the First Approved Small Molecule Splicing Modifier Drug as a Blueprint for Future Transformative Medicines. ACS Med Chem Lett 2021, 12 (6), 874–877. DOI: 10.1021/acsmedchemlett.0c00659 From NLM.

(5) Germer, K.; Leonard, M.; Zhang, X. RNA aptamers and their therapeutic and diagnostic applications. Int J Biochem Mol Biol 2013, 4 (1), 27–40. From NLM.

(6) Kaur, H.; Bruno, J. G.; Kumar, A.; Sharma, T. K. Aptamers in the Therapeutics and Diagnostics Pipelines. Theranostics 2018, 8 (15), 4016–4032. DOI: 10.7150/thno.25958 From NLM.

(7) Menichelli, E.; Lam, B. J.; Wang, Y.; Wang, V. S.; Shaffer, J.; Tjhung, K. F.; Bursulaya, B.; Nguyen, T. N.; Vo, T.; Alper, P. B.;, et al. Discovery of small molecules that target a tertiary-structured RNA. Proc Natl Acad Sci U S A 2022, 119 (48), e2213117119. DOI: 10.1073/pnas.2213117119 From NLM.

(8) Childs-Disney, J. L.; Yang, X.; Gibaut, Q. M. R.; Tong, Y.; Batey, R. T.; Disney, M. D. Targeting RNA structures with small molecules. Nat Rev Drug Discov 2022, 21 (10), 736–762. DOI: 10.1038/s41573-022-00521-4.

(9) Rizvi, N. F.; Howe, J. A.; Nahvi, A.; Klein, D. J.; Fischmann, T. O.; Kim, H.-Y.; McCoy, M. A.; Walker, S. S.; Hruza, A.; Richards, M. P.;, et al. Discovery of Selective RNA-Binding Small Molecules by Affinity-Selection Mass Spectrometry. ACS Chem Biol 2018, 13 (3), 820–831. DOI: 10.1021/acschembio.7b01013.

(10) Rizvi, N. F.; Nickbarg, E. B. RNA-ALIS: Methodology for screening soluble RNAs as small molecule targets using ALIS affinity-selection mass spectrometry. Methods 2019, 167, 28–38. DOI: 10.1016/j.ymeth.2019.04.024.

(11) Wicks, S. L.; Hargrove, A. E. Fluorescent indicator displacement assays to identify and characterize small molecule interactions with RNA. Methods 2019, 167, 3–14. DOI: 10.1016/j.ymeth.2019.04.018.

(12) Zhang, J.; Umemoto, S.; Nakatani, K. Fluorescent Indicator Displacement Assay for Ligand−RNA Interactions. J Am Chem Soc 2010, 132 (11), 3660–3661. DOI: 10.1021/ja100089u.

(13) Connelly, C. M.; Abulwerdi, F. A.; Schneekloth, J. S. Discovery of RNA Binding Small Molecules Using Small Molecule Microarrays. In Small Molecule Microarrays: Methods and Protocols, Uttamchandani, M., Yao, S. Q. Eds.; Springer New York, 2017; pp 157–175.

(14) Benhamou, R. I.; Suresh, B. M.; Tong, Y.; Cochrane, W. G.; Cavett, V.; Vezina-Dawod, S.; Abegg, D.; Childs-Disney, J. L.; Adibekian, A.; Paegel, B. M.;, et al. DNA-encoded library versus RNA-encoded library selection enables design of an oncogenic noncoding RNA inhibitor. Proc Natl Acad Sci U S A 2022, 119 (6), e2114971119. DOI: doi:10.1073/pnas.2114971119.

(15) Shino, A.; Otsu, M.; Imai, K.; Fukuzawa, K.; Morishita, E. C. Probing RNA–Small Molecule Interactions Using Biophysical and Computational Approaches. ACS Chem Biol 2023, 18 (11), 2368–2376. DOI: 10.1021/acschembio.3c00287.

(16) Bennett, J. L.; Nguyen, G. T. H.; Donald, W. A. Protein–Small Molecule Interactions in Native Mass Spectrometry. Chem Rev 2022, 122 (8), 7327–7385. DOI: 10.1021/acs.chemrev.1c00293.

(17) Drinkwater, N.; Vu, H.; Lovell, Kimberly M.; Criscione, Kevin R.; Collins, Brett M.; Prisinzano, Thomas E.; Poulsen, S.-A.; McLeish, Michael J.; Grunewald, Gary L.; Martin, Jennifer L. Fragment-based screening by X-ray crystallography, MS and isothermal titration calorimetry to identify PNMT (phenylethanolamine N-methyltransferase) inhibitors. Biochem J 2010, 431 (1), 51–61. DOI: 10.1042/bj20100651 (acccessed 9/3/2024).

(18) Woods, L. A.; Dolezal, O.; Ren, B.; Ryan, J. H.; Peat, T. S.; Poulsen, S.-A. Native State Mass Spectrometry, Surface Plasmon Resonance, and X-ray Crystallography Correlate Strongly as a Fragment Screening Combination. J Med Chem 2016, 59 (5), 2192–2204. DOI: 10.1021/acs.jmedchem.5b01940.

(19) Largy, E.; König, A.; Ghosh, A.; Ghosh, D.; Benabou, S.; Rosu, F.; Gabelica, V. Mass Spectrometry of Nucleic Acid Noncovalent Complexes. Chem Rev 2022, 122 (8), 7720–7839. DOI: 10.1021/acs.chemrev.1c00386.

(20) Seth, P. P.; Miyaji, A.; Jefferson, E. A.; Sannes-Lowery, K. A.; Osgood, S. A.; Propp, S. S.; Ranken, R.; Massire, C.; Sampath, R.; Ecker, D. J.;, et al. SAR by MS: Discovery of a New Class of RNA-Binding Small Molecules for the Hepatitis C Virus: Internal Ribosome Entry Site IIA Subdomain. J Med Chem 2005, 48 (23), 7099–7102. DOI: 10.1021/jm050815o.

(21) Wolff, P.; Ennifar, E. Native Electrospray Ionization Mass Spectrometry of RNA-Ligand Complexes. In RNA Spectroscopy: Methods and Protocols, Arluison, V., Wien, F. Eds.; Springer US, 2020; pp 111–118.

(22) Tuerk, C.; Gold, L. Systematic evolution of ligands by exponential enrichment: RNA ligands to bacteriophage T4 DNA polymerase. Science 1990, 249 (4968), 505–510. DOI: 10.1126/science.2200121 From NLM.

(23) Krause, K. M.; Serio, A. W.; Kane, T. R.; Connolly, L. E. Aminoglycosides: An Overview. Cold Spring Harb Perspect Med 2016, 6 (6). DOI: 10.1101/cshperspect.a027029 From NLM.

(24) Hermann, T. Chemical and functional diversity of small molecule ligands for RNA. Biopolymers 2003, 70 (1), 4–18. DOI: 10.1002/bip.10410.

(25) Kelly, M. L.; Chu, C. C.; Shi, H.; Ganser, L. R.; Bogerd, H. P.; Huynh, K.; Hou, Y.; Cullen, B. R.; Al-Hashimi, H. M. Understanding the characteristics of nonspecific binding of drug-like compounds to canonical stem-loop RNAs and their implications for functional cellular assays. RNA 2021, 27 (1), 12–26. DOI: 10.1261/rna.076257.120 From NLM.

(26) Keller, K. M.; Breeden, M. M.; Zhang, J.; Ellington, A. D.; Brodbelt, J. S. Electrospray ionization of nucleic acid aptamer/small molecule complexes for screening aptamer selectivity. J Mass Spectrom 2005, 40 (10), 1327–1337. DOI: 10.1002/jms.915.

(27) Lipinski, C. A.; Lombardo, F.; Dominy, B. W.; Feeney, P. J. Experimental and computational approaches to estimate solubility and permeability in drug discovery and development settings1PII of original article: S0169-409X(96)00423-1. The article was originally published in Advanced Drug Delivery Reviews 23 (1997) 3–25.1. *Adv Drug Deliv Rev* 2001, *46* (1), 3-26. DOI: 10.1016/S0169-409X(00)00129-0.

(28) Suresh, B. M.; Taghavi, A.; Childs-Disney, J. L.; Disney, M. D. Fragment-Based Approaches to Identify RNA Binders. J Med Chem 2023, 66 (10), 6523–6541. DOI: 10.1021/acs.jmedchem.3c00034.

(29) Lee, M.-K.; Bottini, A.; Kim, M.; Bardaro, M. F.; Zhang, Z.; Pellecchia, M.; Choi, B.-S.; Varani, G. A novel small-molecule binds to the influenza A virus RNA promoter and inhibits viral replication. Chem Commun 2014, 50 (3), 368–370, 10.1039/C3CC46973E. DOI: 10.1039/C3CC46973E.

(30) Lundquist, K. P.; Panchal, V.; Gotfredsen, C. H.; Brenk, R.; Clausen, M. H. Fragment-Based Drug Discovery for RNA Targets. ChemMedChem 2021, 16 (17), 2588–2603. DOI: 10.1002/cmdc.202100324.

(31) Wang, Y.; Rando, R. R. Specific binding of aminoglycoside antibiotics to RNA. Chem & Biol 1995, 2 (5), 281–290. DOI: 10.1016/1074-5521(95)90047-0.

(32) Jiang, L.; Patel, D. J. Solution structure of the tobramycin–RNA aptamer complex. Nat Struct Biol 1998, 5 (9), 769–774. DOI: 10.1038/1804.

(33) Kwon, M.; Chun, S.-M.; Jeong, S.; Yu, J. In Vitro Selection of RNA against Kanamycin B. Molecules and Cells 2001, 11 (3), 303–311. DOI: 10.1016/S1016-8478(23)17040-3.

(34) Kiga, D.; Futamura, Y.; Sakamoto, K.; Yokoyama, S. An RNA aptamer to the xanthine/guanine base with a distinctive mode of purine recognition. Nucleic Acids Res 1998, 26 (7), 1755–1760. DOI: 10.1093/nar/26.7.1755 From NLM.

(35) Zimmermann, G.; Wick, C.; Shields, T.; Jenison, R.; Pardi, A. Molecular interactions and metal binding in the theophylline-binding core of an RNA aptamer. RNA 2000, 6, 659–667.

(36) Jenison, R. D.; Gill, S. C.; Pardi, A.; Polisky, B. High-Resolution Molecular Discrimination by RNA. Science 1994, 263 (5152), 1425–1429. DOI: doi:10.1126/science.7510417.

(37) Lipfert, J.; Doniach, S.; Das, R.; Herschlag, D. Understanding Nucleic Acid–Ion Interactions. Ann Rev Biochem 2014, 83 (Volume 83, 2014), 813–841. DOI: 10.1146/annurev-biochem-060409-092720.

(38) Konermann, L. Addressing a Common Misconception: Ammonium Acetate as Neutral pH “Buffer” for Native Electrospray Mass Spectrometry. J Am Soc Mass Spectrom 2017, 28 (9), 1827–1835. DOI: 10.1007/s13361-017-1739-3.

(39) Konermann, L.; Liu, Z.; Haidar, Y.; Willans, M. J.; Bainbridge, N. A. On the Chemistry of Aqueous Ammonium Acetate Droplets during Native Electrospray Ionization Mass Spectrometry. Anal Chem 2023, 95 (37), 13957–13966. DOI: 10.1021/acs.analchem.3c02546.

(40) Piccolop, S. Biophysical characterization of aptamer-ligand interactions by native mass spectrometry. Univesité de Bordeaux 2019.

(41) Wang, Y.; Killian, J.; Hamasaki, K.; Rando, R. R. RNA Molecules That Specifically and Stoichiometrically Bind Aminoglycoside Antibiotics with High Affinities. Biochem 1996, 35 (38), 12338–12346. DOI: 10.1021/bi960878w.

(42) Kenderdine, T.; Xia, Z.; Williams, E. R.; Fabris, D. Submicrometer Nanospray Emitters Provide New Insights into the Mechanism of Cation Adduction to Anionic Oligonucleotides. Anal Chem 2018, 90 (22), 13541–13548. DOI: 10.1021/acs.analchem.8b03632.

(43) Sterling, H. J.; Prell, J. S.; Cassou, C. A.; Williams, E. R. Protein Conformation and Supercharging with DMSO from Aqueous Solution. J Am Soc Mass Spectrom 2011, 22 (7). DOI: 10.1007/s13361-011-0116-x.

(44) Kulik, M.; Goral, A. M.; Jasiński, M.; Dominiak, P. M.; Trylska, J. Electrostatic interactions in aminoglycoside-RNA complexes. Biophys J 2015, 108 (3), 655–665. DOI: 10.1016/j.bpj.2014.12.020 From NLM.

(45) Disney, M. D.; Winkelsas, A. M.; Velagapudi, S. P.; Southern, M.; Fallahi, M.; Childs-Disney, J. L. Inforna 2.0: A Platform for the Sequence-Based Design of Small Molecules Targeting Structured RNAs. ACS Chem Biol 2016, 11 (6), 1720–1728. DOI: 10.1021/acschembio.6b00001.

(46) Donlic, A.; Swanson, E. G.; Chiu, L.-Y.; Wicks, S. L.; Juru, A. U.; Cai, Z.; Kassam, K.; Laudeman, C.; Sanaba, B. G.; Sugarman, A.;, et al. R-BIND 2.0: An Updated Database of Bioactive RNA-Targeting Small Molecules and Associated RNA Secondary Structures. ACS Chem Biol 2022, 17 (6), 1556–1566. DOI: 10.1021/acschembio.2c00224.

(47) Morgan, B. S.; Sanaba, B. G.; Donlic, A.; Karloff, D. B.; Forte, J. E.; Zhang, Y.; Hargrove, A. E. R-BIND: An Interactive Database for Exploring and Developing RNA-Targeted Chemical Probes. ACS Chem Biol 2019, 14 (12), 2691–2700. DOI: 10.1021/acschembio.9b00631.

(48) Simpson, M.; Poulsen, S. A. An overview of Australia’s compound management facility: the Queensland Compound Library. ACS Chem Biol 2014, 9 (1), 28–33. DOI: 10.1021/cb400912x From NLM.

(49) Smart Compounds, Compounds Australia Structure Portal. https://smartcompounds.com.au/.

(50) Erlanson, D. A. Introduction to fragment-based drug discovery. Top Curr Chem 2012, 317, 1–32. DOI: 10.1007/128_2011_180 From NLM.

(51) Sander, T.; Freyss, J.; von Korff, M.; Rufener, C. DataWarrior: an open-source program for chemistry aware data visualization and analysis. J Chem Inf Model 2015, 55 (2), 460–473. DOI: 10.1021/ci500588j From NLM.

(52) Marty, M. T.; Baldwin, A. J.; Marklund, E. G.; Hochberg, G. K. A.; Benesch, J. L. P.; Robinson, C. V. Bayesian Deconvolution of Mass and Ion Mobility Spectra: From Binary Interactions to Polydisperse Ensembles. Anal Chem 2015, 87 (8), 4370–4376. DOI: 10.1021/acs.analchem.5b00140.

(53) Yu, Y.; Sternicki, L. M.; Hilko, D. H.; Jarrott, R. J.; Di Trapani, G.; Tonissen, K. F.; Poulsen, S.-A. Investigating Active Site Binding of Ligands to High and Low Activity Carbonic Anhydrase Enzymes Using Native Mass Spectrometry. J Med Chem 2024, 67 (17), 15862–15872. DOI: 10.1021/acs.jmedchem.4c01512.

