## Supplementary Information for "Assessment of native mass spectrometry as a screening method to identify and characterize RNA-targeting small molecules"

### Table of Contents

**Table S1.** Supplier and product numbers for purchased compounds tested for aptamer binding. ................S3

**Figure S1.** Full nMS spectra for tobramycin **1** dose-response binding to the tobramycin RNA aptamer (2 µM, 250 mM $NH_4OAc$, 1% DMSO pH 6.5) without $Mg^{2+}$ present. ..........................................................S4

**Figure S2.** nMS analysis of dose-response binding of the cognate ligands tobramycin **1** and kanamycin B **2** to their respective aptamers (a) tobramycin RNA aptamer, and (b) kanamycin B RNA aptamer (2 µM, 250 mM $NH_4OAc$, 1% DMSO pH 6.5) in the presence of 20 µM $Mg^{2+}$ (as the $Mg(OAc)_2$ salt). ........................S5

**Figure S3.** Full nMS spectra for kanamycin B **2** dose-response binding to the kanamycin B RNA aptamer (2 µM, 250 mM $NH_4OAc$, 1% DMSO pH 6.5) without $Mg^{2+}$ present. ..........................................................S6

**Figure S4.** nMS analysis of the binding of the panel of aminoglycosides **1**, **2**, **5-8** to the (a) tobramycin RNA aptamer, and (b) kanamycin B RNA aptamer (2 µM, 250 mM $NH_4OAc$, 1% DMSO, pH 6.5) in the presence of 20 µM $Mg^{2+}$ (as the $Mg(OAc)_2$ salt). ..........................................................................................S7

**Figure S5.** nMS analysis of the binding of the negative control compounds theophylline **4**, D -glucosamine **9**, caffeine **10** to the tobramycin RNA aptamer (2 µM, 250 mM $NH_4OAc$, 1% DMSO pH 6.5), both in the (a) absence and (b) presence of 20 µM $Mg^{2+}$ (as the $Mg(OAc)_2$ salt). ........................................................S8

**Figure S6.** nMS analysis of the binding of the negative control compounds theophylline **4**, D-glucosamine **9**, caffeine **10** to the kanamycin B RNA aptamer (2 µM, 250 mM $NH_4OAc$, 1% DMSO pH 6.5), both in the (a) absence and (b) presence of 20 µM $Mg^{2+}$ (as the $Mg(OAc)_2$ salt). ................................................................ S9

**Figure S7.** Compound library annotated with compound class, compound name, chemical structure and code number. ................................................................ S10

**Figure S8.** Distribution of chemical properties for screening hits plotted against % binding to each aptamer. ................................................................ S15

**Table S1.** Supplier and product numbers for purchased compounds tested for aptamer binding.

| Compound | Ligand | Supplier | Product Number |
|---|---|---|---|
| **1** | tobramycin | AK Scientific | J10405 |
| **2** | kanamycin B | AK Scientific | J10862 |
| **3** | xanthine | AK Scientific | K177 |
| **4** | theophylline | AK Scientific | J40064 |
| **5** | kanamycin A | AK Scientific | J10752 |
| **6** | neomycin sulfate | AK Scientific | J10828 |
| **7** | paromomycin sulfate | AK Scientific | H914 |
| **8** | gentamicin sulfate | AK Scientific | J95790 |
| **9** | D-glucosamine | Sigma Aldrich | G4875 |
| **10** | caffeine | AK Scientific | J11627 |
| **11** | amikacin disulfate | AK Scientific | J10297 |
| **12** | apramycin sulfate salt | AK Scientific | J10972 |
| **13** | dibekacin sulfate | AK Scientific | H670 |
| **14** | geneticin disulfate | AK Scientific | J97653 |
| **15** | netilmicin sulfate | AK Scientific | J62396 |
| **16** | ribostamycin sulfate salt | AK Scientific | J51266 |
| **17** | streptomycin sulfate | AK Scientific | J96098 |

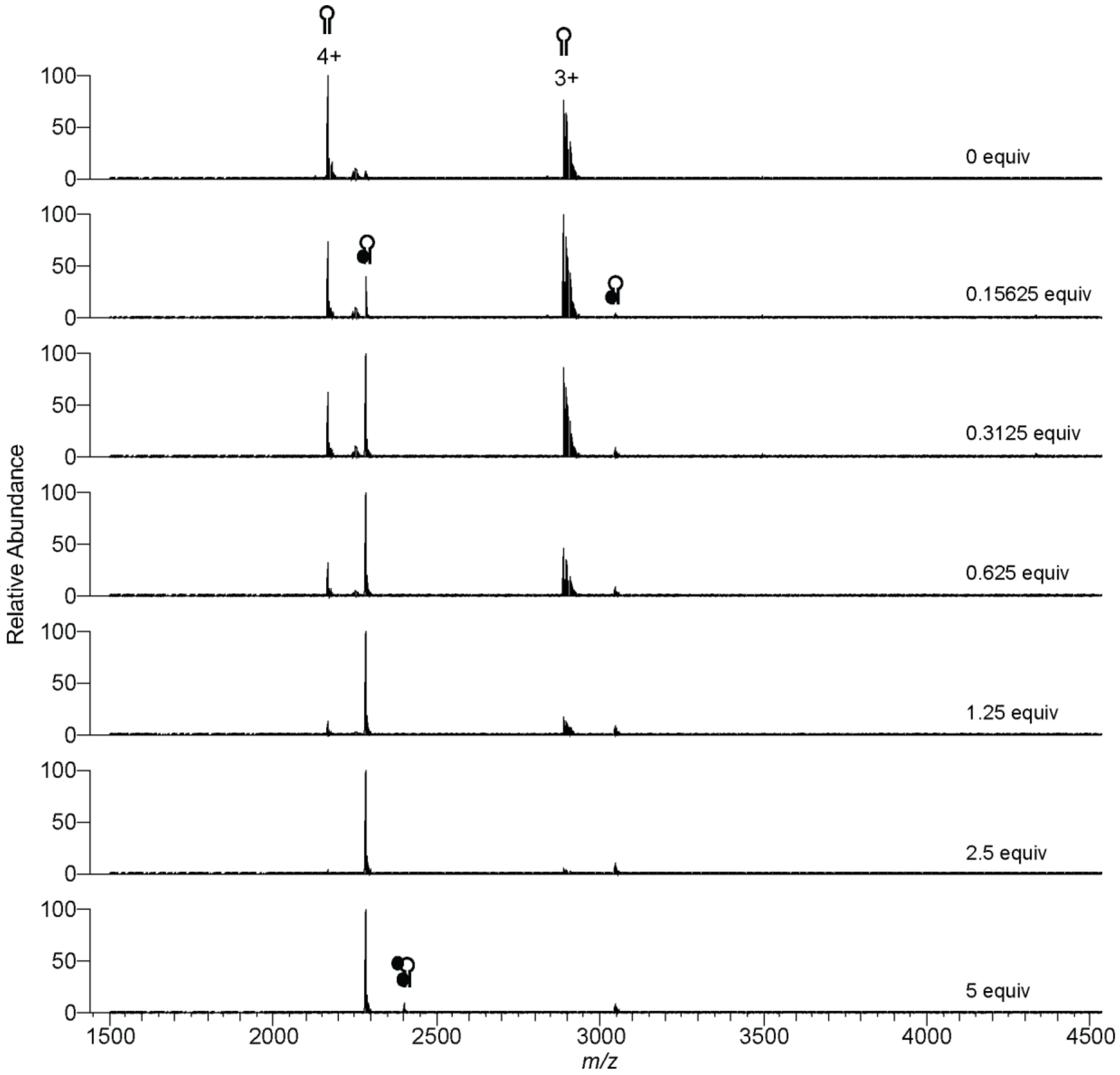


**Figure S1.** Full nMS spectra for tobramycin **1** dose-response binding to the tobramycin RNA aptamer (2 µM, 250 mM $NH_4OAc$, 1% DMSO pH 6.5) without $Mg^{2+}$ present. Hairpin represents aptamer, black circle represents bound ligand **1**. nMS spectra representative of n=2.

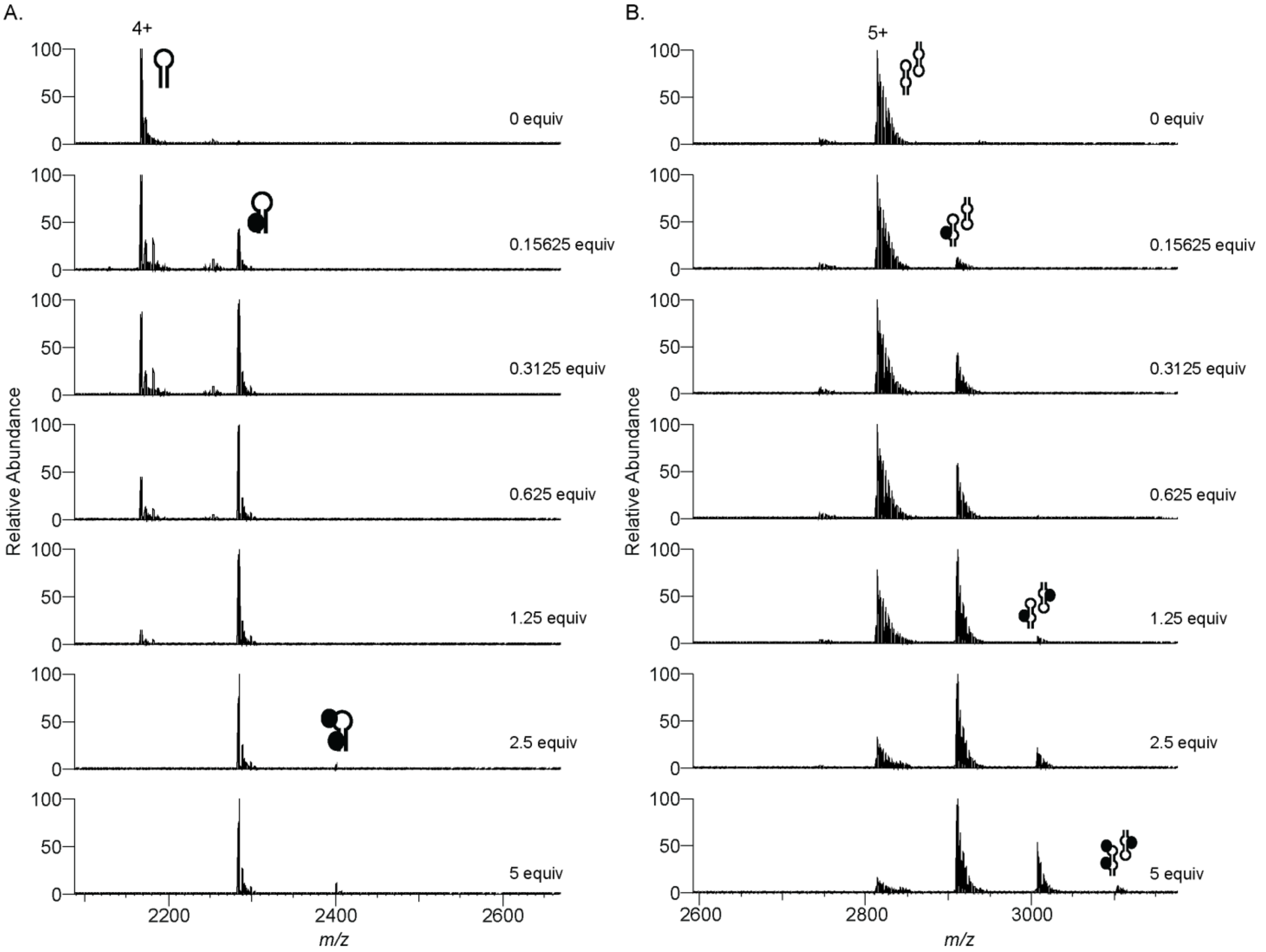


**Figure S2.** nMS analysis of dose-response binding of the cognate ligands tobramycin **1** and kanamycin B **2** to their respective aptamers (a) tobramycin RNA aptamer, and (b) kanamycin B RNA aptamer (2 µM, 250 mM $NH_4OAc$, 1% DMSO pH 6.5) in the presence of 20 µM $Mg^{2+}$ (as the $Mg(OAc)_2$ salt). Spectra for the major observed charge state with ligand binding are shown. Hairpin represents aptamer, double hairpin represents dimeric aptamer, black circle represents bound ligand **1** (a) or **2** (b). nMS spectra representative of n=2.

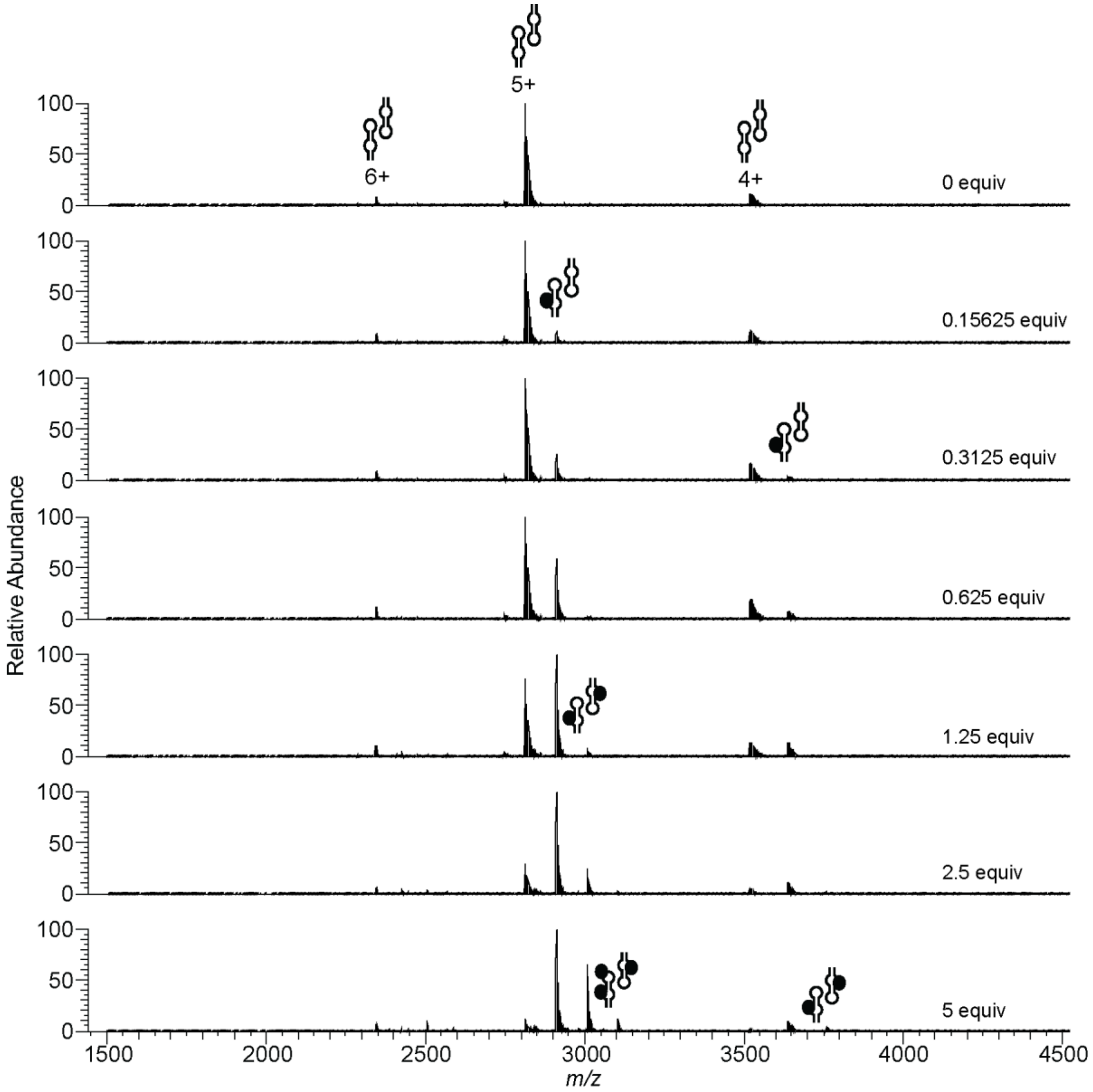


**Figure S3.** Full nMS spectra for kanamycin B **2** dose-response binding to the kanamycin B RNA aptamer (2 µM, 250 mM $NH_4OAc$, 1% DMSO pH 6.5) without $Mg^{2+}$ present. Double hairpin represents dimeric aptamer, black circle represents bound ligand **2**. nMS spectra representative of n=2.

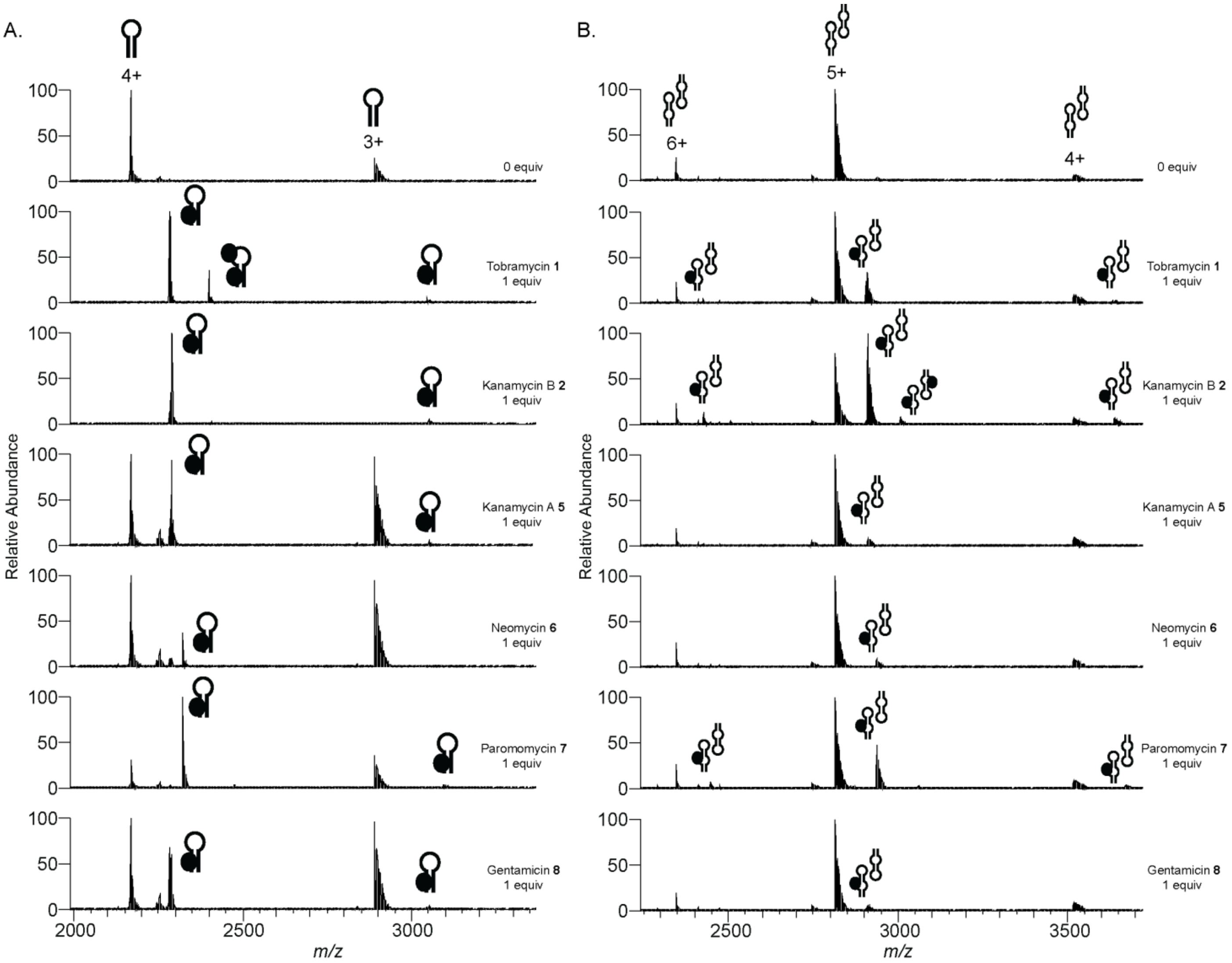


**Figure S4.** nMS analysis of the binding of the panel of aminoglycosides **1**, **2**, **5-8** to the (a) tobramycin RNA aptamer, and (b) kanamycin B RNA aptamer (2 µM, 250 mM $NH_4OAc$, 1% DMSO, pH 6.5) in the presence of 20 µM $Mg^{2+}$ (as the $Mg(OAc)_2$ salt). Hairpin represents aptamer, double hairpin represents dimeric aptamer, black circle represents bound ligands **1**, **2**, **5-8**. nMS spectra representative of n=2.

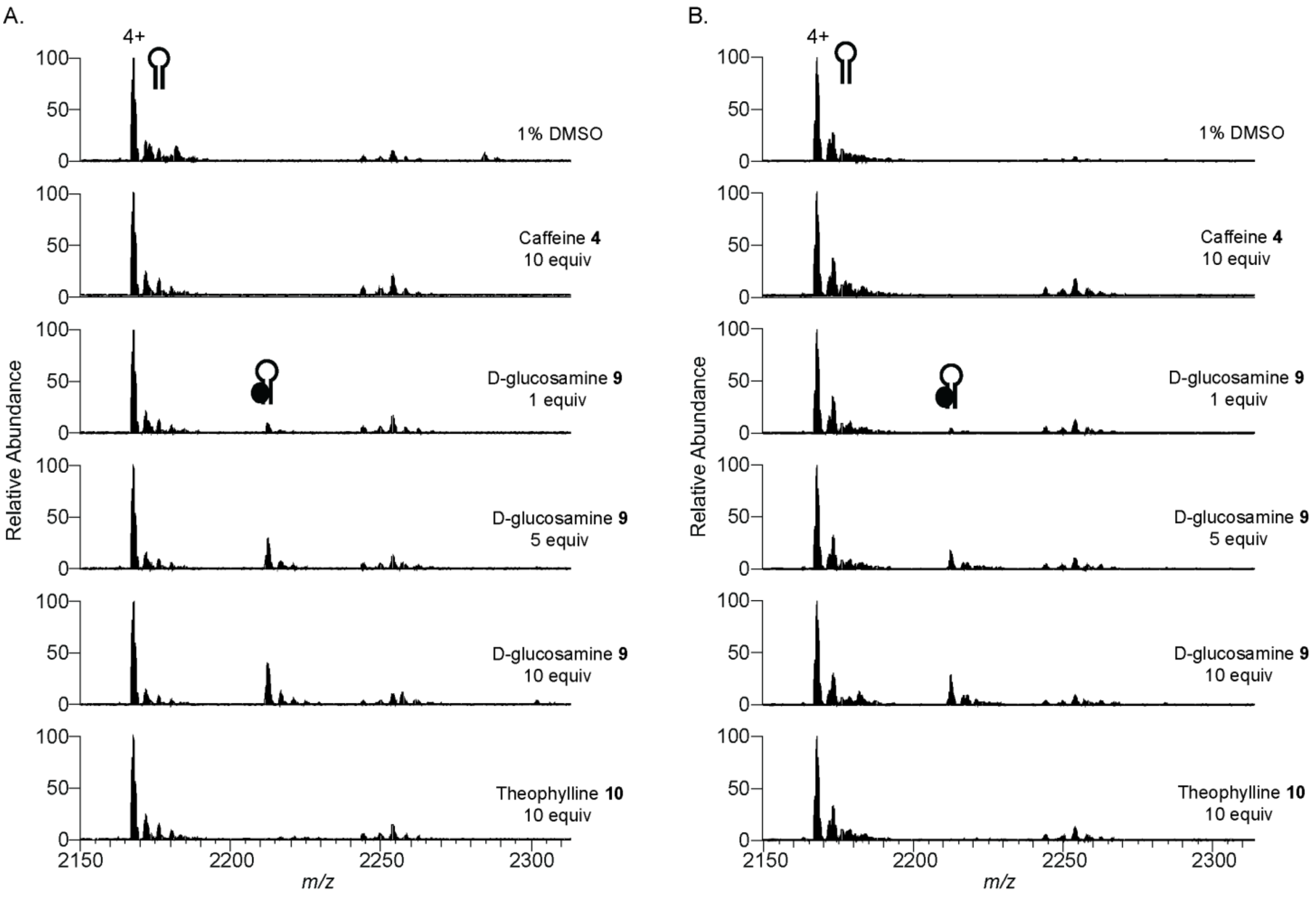


**Figure S5.** nMS analysis of the binding of the negative control compounds theophylline **4**, D -glucosamine **9**, caffeine **10** to the tobramycin RNA aptamer (2 µM, 250 mM $NH_4OAc$, 1% DMSO pH 6.5), both in the (a) absence and (b) presence of 20 µM $Mg^{2+}$ (as the $Mg(OAc)_2$ salt). Spectra for the major observed charge state with ligand binding are shown. Hairpin represents aptamer, black circle represents bound ligand **4**, **9** or **10**. nMS spectra representative of n=2.

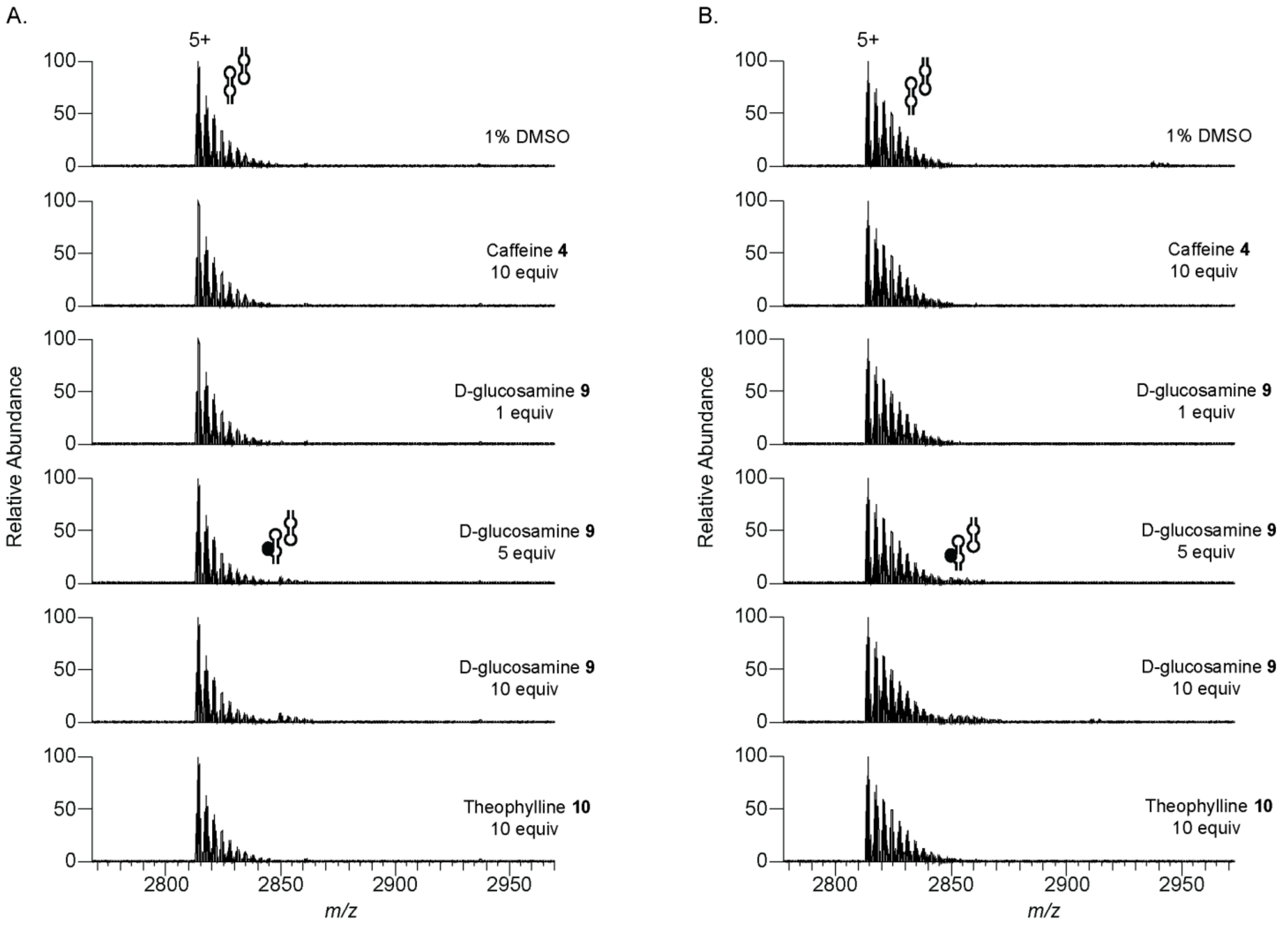


**Figure S6.** nMS analysis of the binding of the negative control compounds theophylline **4**, D-glucosamine **9**, caffeine **10** to the kanamycin B RNA aptamer (2 µM, 250 mM $NH_4OAc$, 1% DMSO pH 6.5), both in the (a) absence and (b) presence of 20 µM $Mg^{2+}$ (as the $Mg(OAc)_2$ salt). Only the predominant charge state observed is shown. Spectra for the major observed charge state with ligand binding are shown. Double hairpin represents dimeric aptamer, black circle represents bound ligand **4**, **9** or **10**. nMS spectra representative of n=2.

**Figure S7.** Compound library annotated with compound class, compound name, chemical structure and code number. The library comprises 47 antibiotic drugs and 30 other drugs. (a) Aminoglycosides **11-17**, (b) tetracyclines **18-27**, (c) anthracyclines **28-34**, (d) macrolides **35-46**, (e) lincosamides **47-48**, (f) fluoroquinolines **49-54**, (g) oxazolodinones **55-57**) (h) drugs of MW >300 Da **58-72** and (i) fragment drugs of MW <300 Da **73-87**.

(a) Aminoglycosides **11-17**

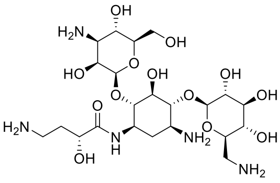
Amikacin, 11

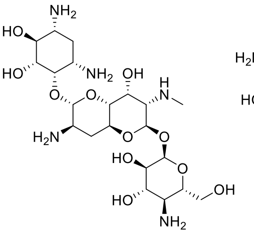
Apramycin, 12

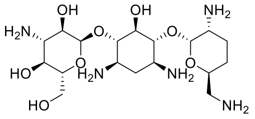
Dibekacin, 13

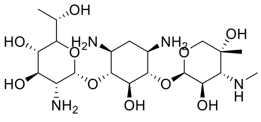
Geneticin, 14

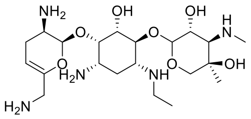
Netilmicin, 15

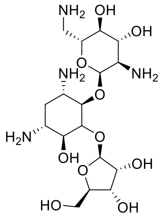
Ribostamycin, 16

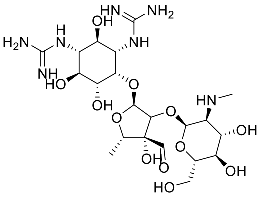
Streptomycin, 17

(b) Tetracyclines **18-27**

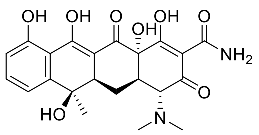
Tetracycline, 18

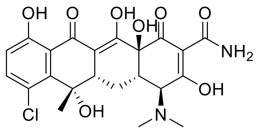
Chlortetracycline, 19

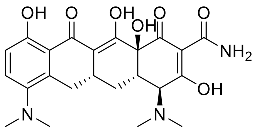
Minocycline, 20

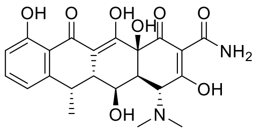
Doxycyclinehyclate, 21

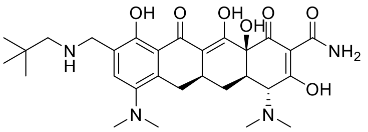
Omadacycline, 22

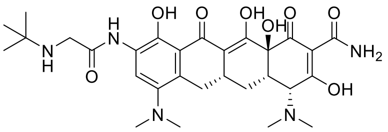
Tigecycline, 23

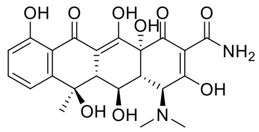
Oxytetracycline, 24

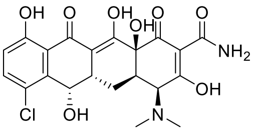
Demeclocycline, 25

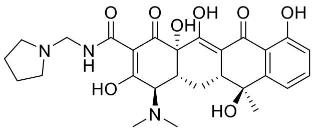
Rolitetracycline, 26

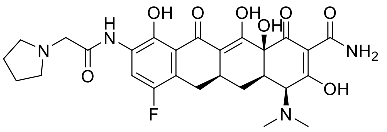
Eravacycline, 27

(c) Anthracyclines **28-34**

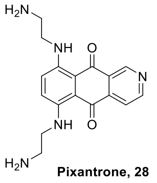

**Pixantrone, 28**

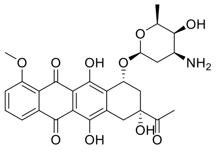

**Daunorubicin, 29**

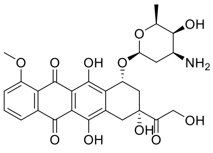

**Doxorubicin, 30**

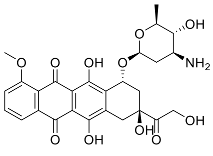

**Epirubicin, 31**

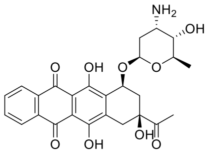

**Idarubicin, 32**

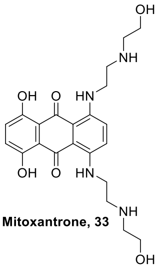

**Mitoxantrone, 33**

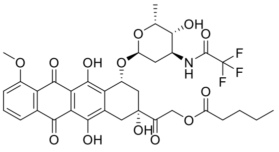

**Valrubicin, 34**

(d) Macrolides **35-46**

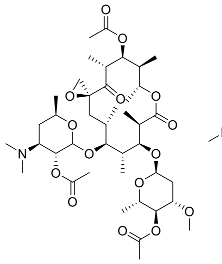

**Troleandomycin, 35**

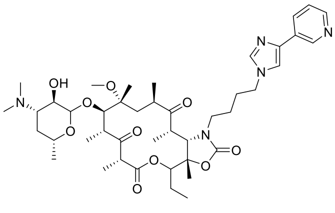

**Telithromycin, 36**

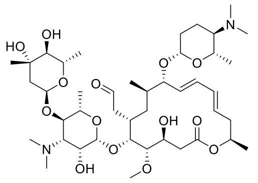

**Spiramycin, 37**

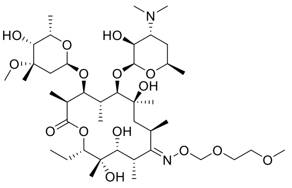

**Roxithromycin, 38**

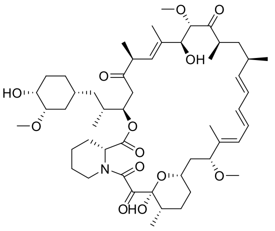

**Rapamycin, 39**

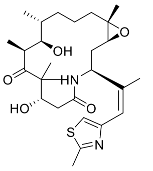

**Ixabepilone, 40**

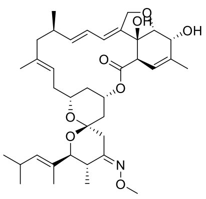

**Moxidectin, 41**

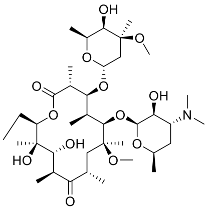

**Clarithromycin, 42**

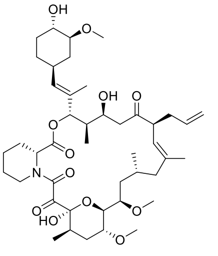

**Tacrolimus,43**

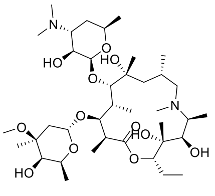

**Azithromycin, 44**

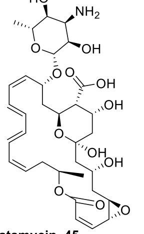

**Natamycin, 45**

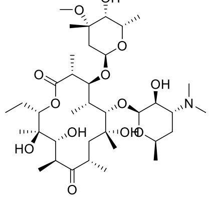

**Erythromycin, 46**

(e) Lincosamides, **47-48**

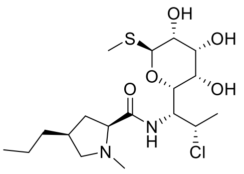

**Clindamycin, 47**

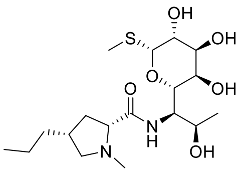

**Lincomycin, 48**

(f) Fluoroquinolines, **49-54**

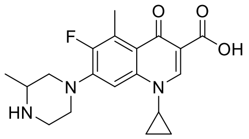

**Grepafloxacin, 49**

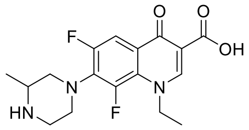

**Lomefloxacin, 50**

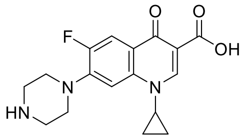

**Ciprofloxacin, 51**

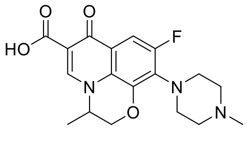

**Ofloxacin, 52**

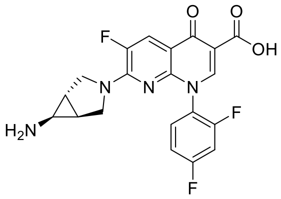

**Trovafloxacin, 53**

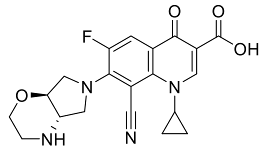

**Finafloxacin, 54**

(g) oxazolodinones **55-57**

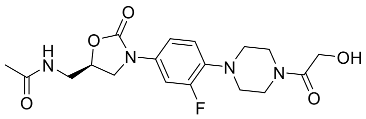

**Eperezolid, 55**

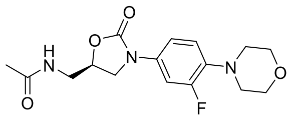

**Linezolid, 56**

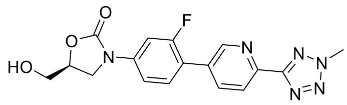

**Tedizolid, 57**

(h) Drug set >300 Da **58-72**

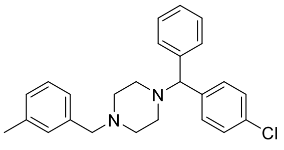

**Meclizine, 58**

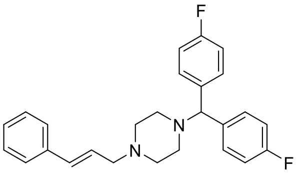

**Flunarizine, 59**

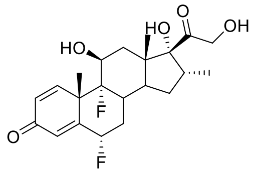

**Flumethasone, 60**

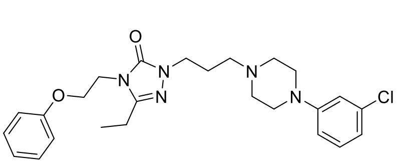

**Nefazodone, 61**

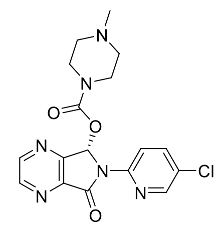

**Eszopiclone, 62**

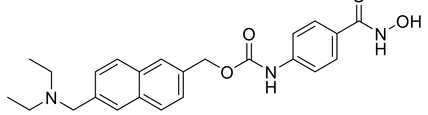

**Givinostat, 63**

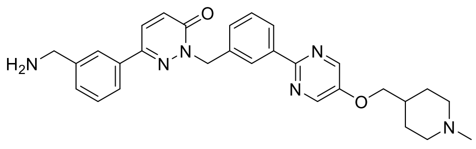

**Tepotinib, 64**

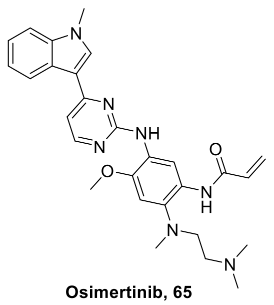

**Osimertinib, 65**

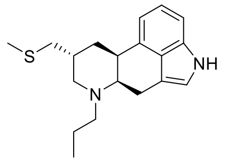

**Pergolide, 66**

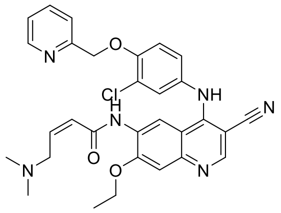

**Neratinib, 67**

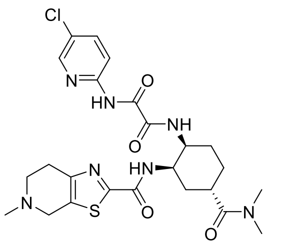

**Edoxaban, 68**

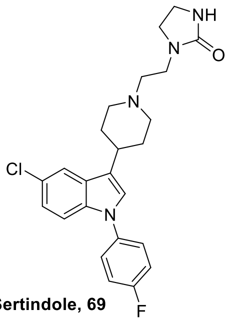

**Sertindole, 69**

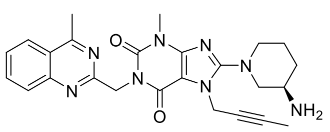

**Linagliptin, 70**

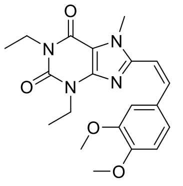

**Istradefylline, 71**

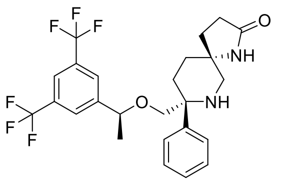

**Rolapitant, 72**

(i) Fragment drug set <300 Da **73-87**

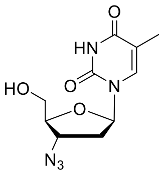

**Zidovudine, 73**

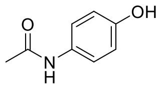

**Acetaminophen, 74**

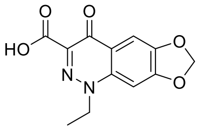

**Cinoxacin, 75**

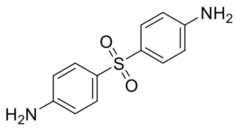

**Dapsone, 76**

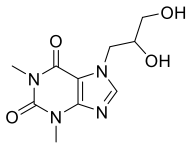

**Dyphylline, 77**

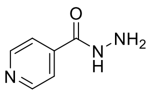

**Isonizid, 78**

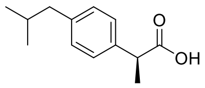

**Dexibuprofen, 79**

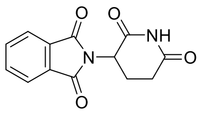

**Thalidomide, 80**

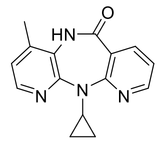

**Nevirapine, 81**

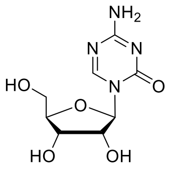

**Azacitidine, 82**

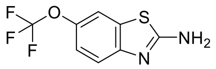

**Riluzole, 83**

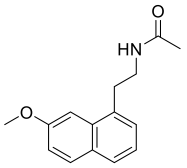

**Agomelatine, 84**

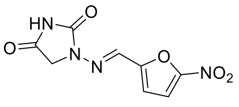

**Nitrofurantoin, 85**

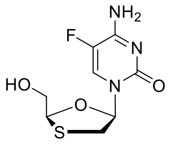

**Emtricitabine, 86**

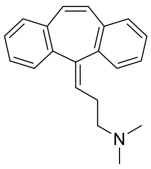

**Cyclobenzaprine, 87**

**Figure S8.** Distribution of chemical properties for screening hits plotted against % binding to each aptamer. (a) MW (Da); (b) number of chiral centres; (c) number of H-bond acceptors; (d) number of H-bond donors; (e) cLogP; (f) PSA; (g) number of rotatable bonds.

(a)

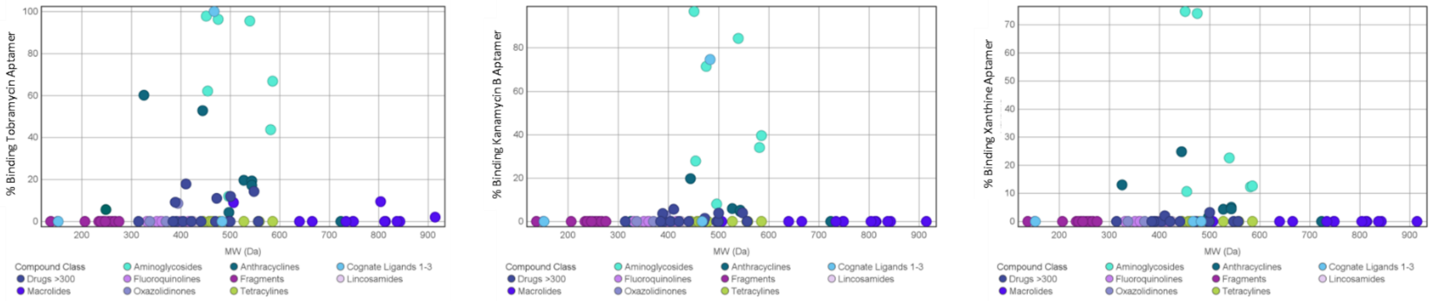


(b)

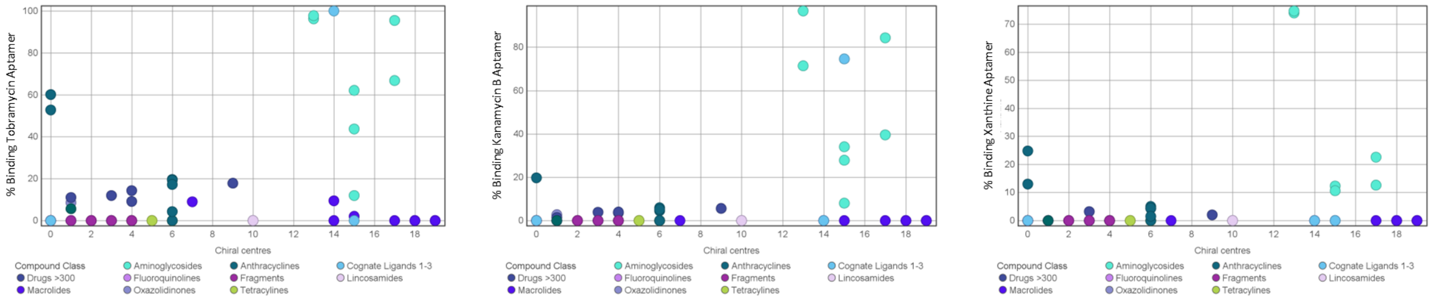


(c)

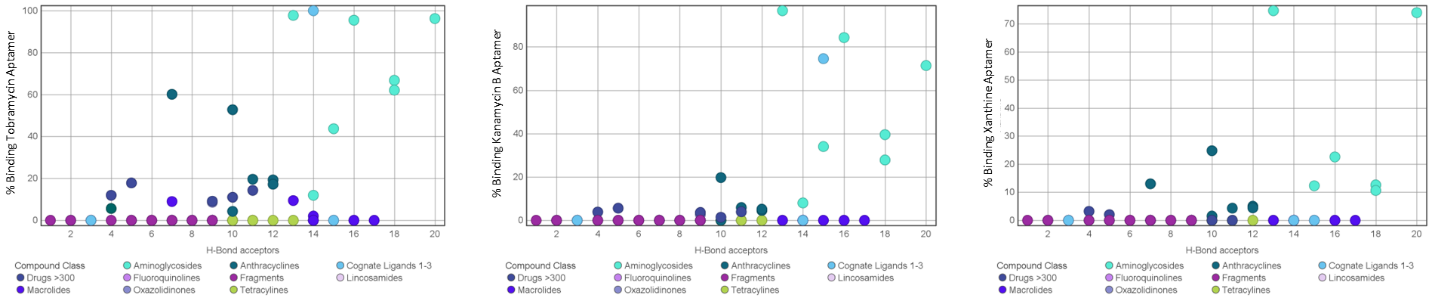


(d)

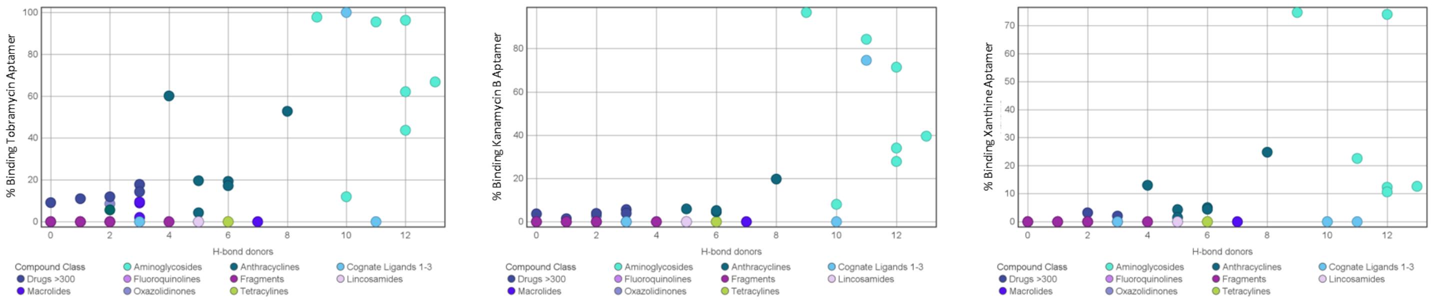

(e)

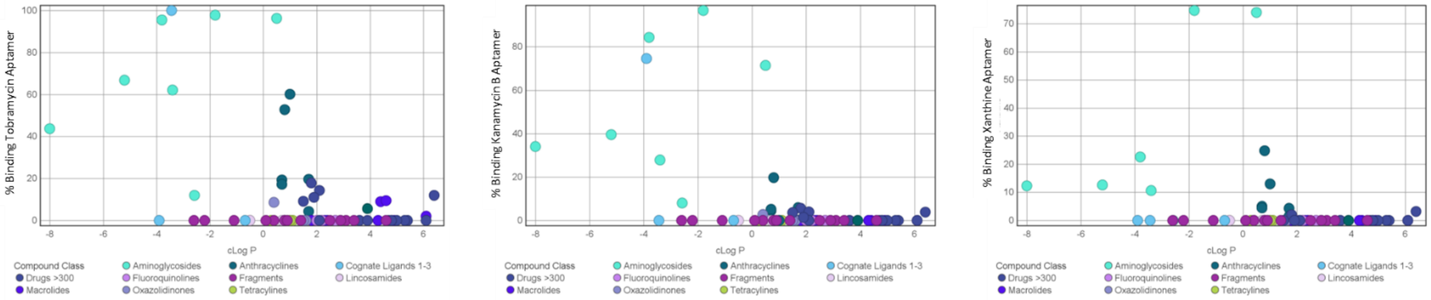
% Binding Tobramycin Aptamer
% Binding Kanamycin B Aptamer
% Binding Xanthine Aptamer
cLog P
Compound Class
Aminoglycosides
Anthracyclines
Cognate Ligands 1-3
Drugs >300
Fluoroquinolines
Fragments
Lincosamides
Macrolides
Oxazolidinones
Tetracylines

(f)

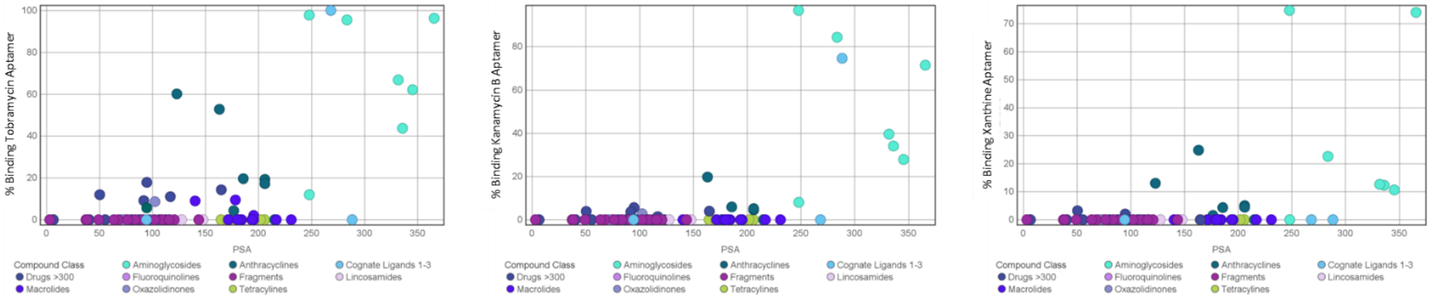
% Binding Tobramycin Aptamer
% Binding Kanamycin B Aptamer
% Binding Xanthine Aptamer
PSA
Compound Class
Aminoglycosides
Anthracyclines
Cognate Ligands 1-3
Drugs >300
Fluoroquinolines
Fragments
Lincosamides
Macrolides
Oxazolidinones
Tetracylines

(g)

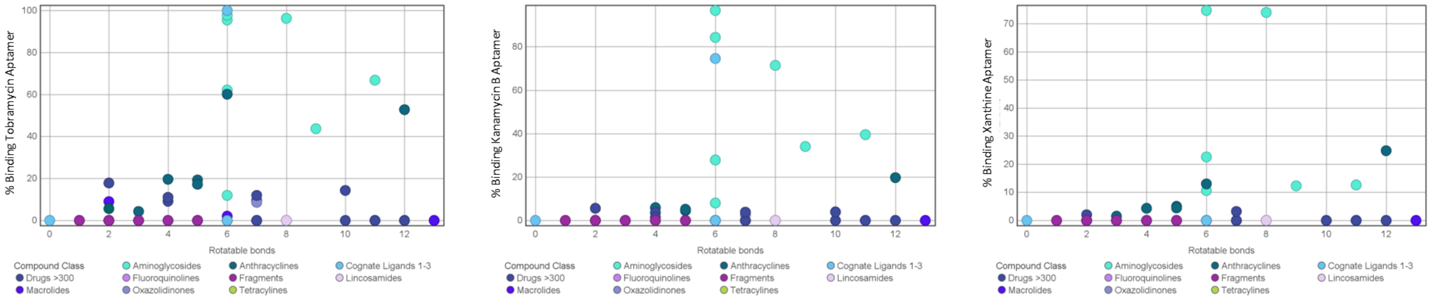
% Binding Tobramycin Aptamer
% Binding Kanamycin B Aptamer
% Binding Xanthine Aptamer
Rotatable bonds
Compound Class
Aminoglycosides
Anthracyclines
Cognate Ligands 1-3
Drugs >300
Fluoroquinolines
Fragments
Lincosamides
Macrolides
Oxazolidinones
Tetracylines
